# Closing and opening of the RNA polymerase trigger loop

**DOI:** 10.1101/817080

**Authors:** Abhishek Mazumder, Miaoxin Lin, Achillefs N. Kapanidis, Richard H. Ebright

## Abstract

The RNA polymerase (RNAP) trigger loop (TL) is a mobile structural element of the RNAP active center that, based on crystal structures, has been proposed to cycle between an “unfolded”/“open” state that allows an NTP substrate to enter the active center and a “folded”/“closed” state that holds the NTP substrate in the active center. Here, by quantifying single-molecule fluorescence resonance energy transfer between a first fluorescent probe in the TL and a second fluorescent probe elsewhere in RNAP or in DNA, we detect and characterize TL closing and opening in solution. We show that the TL closes and opens on the millisecond timescale; we show that TL closing and opening provides a checkpoint for NTP complementarity, NTP ribo/deoxyribo identity, and NTP tri/di/monophosphate identity, and serves as a target for inhibitors; and we show that one cycle of TL closing and opening typically occurs in each nucleotide addition cycle in transcription elongation.

## Introduction

Transcription, the first and most highly regulated process in gene expression, entails transcription initiation, in which RNA polymerase (RNAP) binds to DNA and begins synthesis of an RNA molecule; followed by transcription elongation, in which RNAP extends the RNA molecule; followed by transcription termination, in which RNAP releases the RNA molecule (*1–3*). During transcription elongation, RNAP uses a “stepping” mechanism, in which RNAP translocates relative to DNA by one base pair for each nucleotide added to the RNA molecule (*2, 4–6*). Each “step” occurs as part of a “nucleotide-addition cycle” comprising: (i) RNAP translocation, (ii) nucleoside-triphosphate (NTP) binding, (iii) phosphodiester-bond formation, and (iv) pyrophosphate release (*2, 5, 7*).

A proposed key player in the nucleotide-addition cycle is the RNAP “trigger loop” (TL), a mobile structural element of the RNAP active center that is conserved in RNAP from bacteria through humans (*2, 5, 8–11*). Crystal structures of transcription elongation complexes indicate that the TL can adopt (i) an “unfolded,” or “open,” TL conformation that allows an NTP to enter the RNAP active center (observed in crystal structures without a bound NTP; *2, 5, 8-11*; Fig. 1a, left); and (ii) a “folded,” or “closed,” TL conformation that holds an NTP in the RNAP active center (observed in crystal structures with a bound NTP; *2, 5, 8-11*; Fig. 1a, right). The open and closed TL conformations observed in crystal structures differ by a large--up to ~20 Å--displacement of residues at the tip of the trigger loop (*2, 5, 8-11*; Fig. 1a-b). It has been hypothesized that the open and closed TL conformations observed in crystal structures occur in solution and are crucial for RNAP function in solution. Specifically, TL closing has been hypothesized to contribute to discrimination between complementary and non-complementary NTPs and to contribute to discrimination between NTPs and dNTPs (*2, 5, 8–28*). TL closing also has been hypothesized to contribute to catalysis of phosphodiester-bond formation, by increasing the order of the RNAP active center, by excluding solvent from the RNAP active center, and by positioning residues to participate in electrostatic, general-acid/general-base, or conformational stabilization of the γ-phosphate/β-phosphate leaving group (*2, 5, 8–28*). It further has been hypothesized that the TL returns to its initial, unfolded, open conformational state on or after phosphodiester-bond formation, thereby re-opening the RNAP active center and permitting pyrophosphate release and RNAP translocation (*2, 5, 8–28*).

**Fig. 1.**
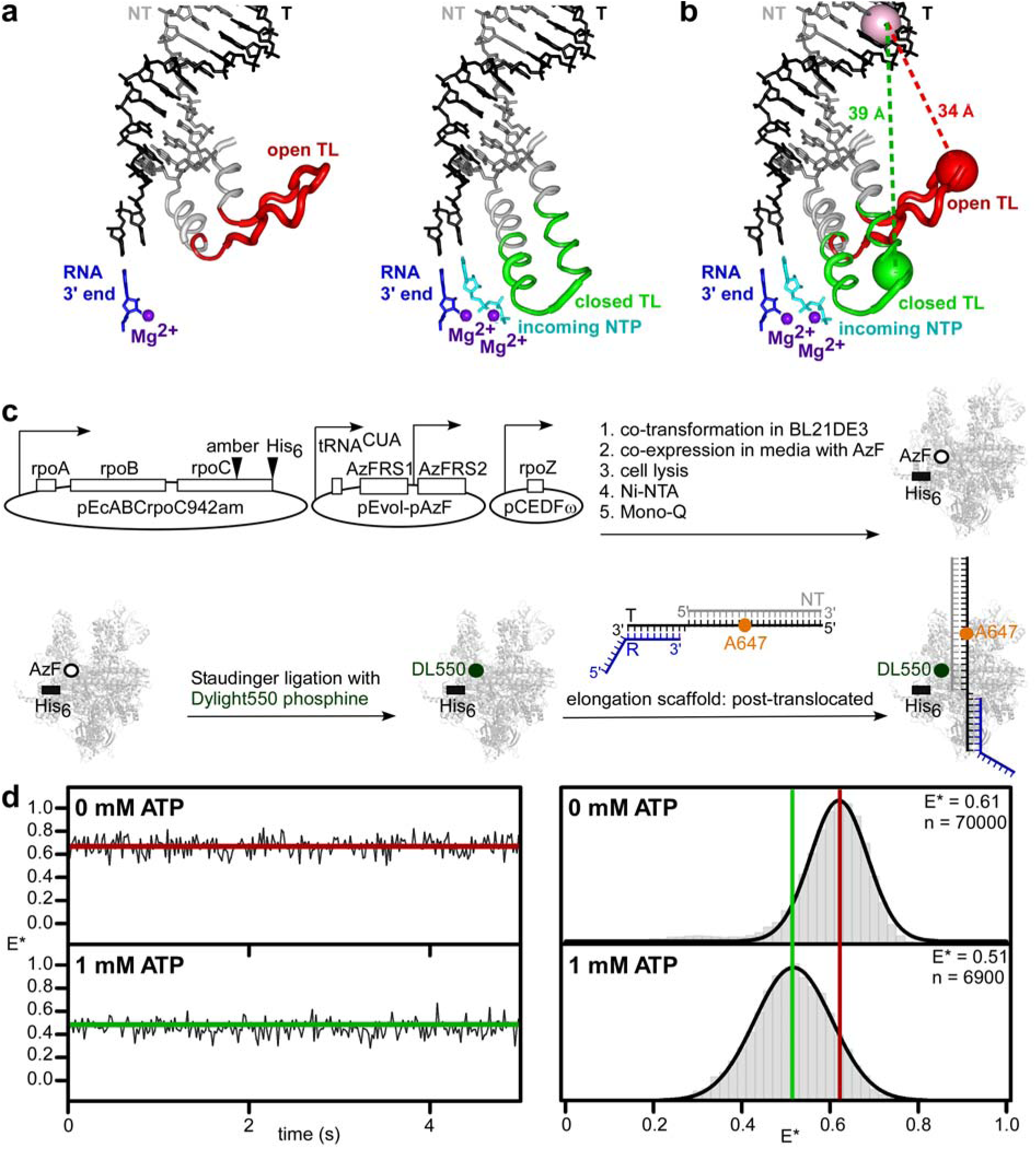
Use of smFRET to detect and characterize TL closing and opening in solution. **a,** Open-TL (left subpanel) and closed-TL (right subpanel) conformational states as observed in crystal structures of *T. thermophilus* RNAP (*9,29*; PDB 1ZYR and PDB 2O5J). Gray and red ribbon, RNAP trigger helices and TL in open-TL state; gray and green ribbon, RNAP trigger helices and TL in closed-TL state, gray and black sticks, DNA non-template and template strands; blue sticks, RNA 3’ nucleotide; cyan stick, incoming NTP; purple sphere in left panel, catalytic Mg^2+^ ions Mg^2+^(I) (left and right subpanels) and Mg^2+^(II) (right subpanel). **b**, Measurement of smFRET between first fluorescent probe incorporated at tip of RNAP TL (red sphere for open-TL state; green sphere for closed-TL) and second fluorescent probe incorporated into downstream DNA (pink sphere). Estimated probe-probe distances are ~34 Å for open-TL state and ~39 Å for closed-TL state. **c,** Use of unnatural--amino-acid mutagenesis (first reaction arrow), Staudinger ligation (second reaction arrow), and TEC reconstitution (third reaction arrow) to prepare sample for measurement of smFRET between first fluorescent probe incorporated at tip of RNAP TL and second fluorescent probe incorporated into downstream DNA (see Methods). Black open ovals, open rectangles, two-segment arrows, arrowhead labelled “amber,” and arrowhead labelled “His_6_” denote plasmids, genes, promoters, amber codon in TL coding sequence in gene for RNAP β’ subunit, and hexahistidine coding sequence at 3’ end of gene for RNAP β’ subunit. Gray ribbon structure, black open circle, green filled circle, and black bar denote RNAP, unnatural amino acid 4-azidophenylalanine in RNAP TL, fluorescent probe Dylight 550 in RNAP TL, and hexahistidine tag at C-terminus of RNAP β’ subunit. Gray lines, black lines, blue lines, and orange filled circle denote nucleic-acid scaffold comprising nontemplate-strand DNA, template-strand DNA, RNA, and fluorescent probe Alexa647. **d,** smFRET data for TEC in post-translocated state in absence of NTP (top) and in presence of saturating concentration of complementary NTP (1 mM ATP; bottom). Left subpanels, representative time traces of donor-acceptor FRET efficiency, E*, showing open-TL (top, red) and closed-TL (bottom, green) states. Right subpanel, histograms and Gaussian fits of E*, showing mean E* values for open-TL (red line) and closed-TL (green line) states; n, number of frames.

According to these proposals, an RNAP active-center conformational cycle, comprising TL closing followed by TL opening, is coupled to each nucleotide-addition cycle (*2, 5, 8–28*). Consistent with these proposals, amino acid substitutions predicted to interfere with TL closing and opening (e.g., substitution of TL Gly residues with conformationally more restricted Ala residues) interfere with nucleotide addition (*14–16, 18–19, 25–27*). Further consistent with these proposals, small molecules predicted to interact with, and trap, the TL in one conformational state interfere with nucleotide addition (9, *13–14, 22, 29–36*). Further consistent with these proposals, a “Cys-pair reporter strategy” detects differences in rates of disulfide-bond formation that correlate with predicted TL closed and open states (*22, 25*). However, no direct observation of TL closed and open states in solution has been reported. In particular, no direct observation of TL closing and opening during real-time, active transcription elongation in solution has been reported, and, as a result, it has not been possible to test directly the hypothesis that an RNAP active-center conformational cycle comprising TL closing followed by TL opening is coupled to each nucleotide-addition cycle.

Here, by use of single-molecule fluorescence resonance energy transfer (smFRET), we directly detect and directly characterize TL closing and opening in solution, including TL closing and opening during real-time, active transcription elongation in solution.

### Use of smFRET to detect and characterize TL closing and opening in solution

In a first set of experiments, we assessed smFRET between a first fluorescent probe, serving as donor, in the RNAP TL and a second fluorescent probe, serving as acceptor, in DNA (Fig. 1b). We used a procedure comprising: (i) incorporation of the fluorescent probe DyLight 550, serving as donor, in the RNAP TL, by use of unnatural-amino-acid mutagenesis and Staudinger ligation (*37–40*); (ii) incorporation of the fluorescent probe Alexa647, serving as acceptor, in the template strand of a nucleic-acid scaffold comprising a template-strand DNA oligonucleotide, a non-template strand DNA oligonucleotide, and a non-extendable, 3’-deoxyribonucleotide-containing RNA oligonucleotide; (iii) assembly and analysis of a doubly labelled transcription elongation complex (TEC) from the resulting labelled RNAP and labelled nucleic-acid scaffold; (iv) immobilization of the doubly labelled TEC, through a hexahistidine tag on RNAP on a surface functionalized with anti-hexahistidine tag-antibody; and (v) measurement of smFRET (Figs. 1c and S1–S2).

We evaluated ten potential labelling sites in the TL of *Escherichia coli* RNAP: i.e., β’ subunit residues 933-942. Seven sites were located in a TL segment that is unfolded in the open-TL state but folded in the closed-TL state (β’ residues 933-939), and three sites were located in a TL segment--the TL tip--that is unfolded in both the open-TL and closed-TL states (β’ residues 940-942). All ten sites were located in the TL region immediately preceding the species-specific sequence insertion present in the TL of *E. coli* RNAP (“SI3”; also referred to as “β’ G non-conserved domain”; β’ residues 943-1130; *41*). All ten sites were sites that exhibit large, ~15-25 Å, differences in Cα positions in crystal structures of the open-TL and closed-TL states (Fig. 1a). For each of the ten sites, we prepared labelled RNAP and then assessed labelling efficiency, labelling specificity, and transcriptional activity. For three sites--β’ residues 940, 941, and 942--the labelled RNAP derivative retained ≥50% of the transcriptional activity of unlabelled wild-type RNAP, and, for one site--β’ residue 942--the labelled RNAP derivative retained ≥90% of the transcriptional activity of unlabelled wild-type RNAP (Fig. S2). We performed subsequent experiments using RNAP labelled at β’ residue 942.

We evaluated two labelling sites in DNA, both located in the double-stranded DNA segment downstream of the RNAP active center: i.e., template-strand position +12 and template-strand position +10 (Fig. 1b). The two labelling sites in DNA were selected to provide the largest, and second-largest predicted differences in probe-probe distance, and the corresponding largest and second-largest predicted differences in smFRET, for the open-TL state vs. the closed-TL state (high FRET in open-TL state; low FRET in closed-TL state; Fig. 1b). The labelling site at position +12 was used in most experiments; the labelling site at position +10 was used in selected additional experiments.

We analyzed TECs in two translocational states: (i) the pre-translocated state (i.e., the state at the start of the nucleotide-addition cycle, with an RNA-DNA hybrid of 10 bp, and with an RNAP active-center addition site--“A site”--occupied by the RNA 3’ end; sequence in Fig. S1b), and (ii) the post-translocated state (i.e., the state in which RNAP has stepped forward by 1 bp, reducing the length of the RNA-DNA hybrid to 9 bp, and rendering the RNAP active-center A site unoccupied and available to bind the next NTP; sequence in Fig. S1a). For analysis of the post-translocated state, we employed a non-extendable, 3’-deoxyribonucleotide-containing RNA in order to allow NTP binding but not NTP addition.

To monitor the distance between fluorescent probes in the resulting donor-acceptor labelled elongation complexes, we used total internal reflection microscopy with alternating laser excitation (TIRF-ALEX) (*37, 39, 40, 42–45*) and quantified smFRET from single TECs immobilized, through a hexahistidine tag on RNAP, on anti-hexahistidine-tag-antibody-functionalized glass cover slips. TIRF-ALEX allows filtering of data to identify only single molecules that contain both a donor and an acceptor, eliminating complications due to incomplete labelling and imperfect reconstitution of TECs (*37, 39, 40, 42–45*). The results provide equilibrium population distributions of apparent smFRET efficiency, E*. Monitoring E* time trajectories of individual TECs reports on the kinetics of TL conformational cycling.

For TECs in the post-translocated state in the absence of an NTP, we observed unimodal E* distributions with mean E* of 0.61, indicative of an open-TL state with a probe-probe mean distance, R, of 49 Å (Fig. 1d; top right). There was no indication of a lower-FRET, closed-TL state within the temporal resolution of the experiment (~20 ms), and there likewise was no indication of a lower-FRET, closed-TL state in experiments employing confocal optical microscopy to provide higher temporal resolution (~1 ms; Fig. S3, right). We conclude that, in a post-translocated TEC state, in the absence of an NTP, the TL is predominantly, potentially exclusively, in an open-TL state.

In contrast, for TECs in the post-translocated state in the presence of a saturating concentration of the complementary NTP (ATP, using a template directing addition of A), the distribution was unimodal, but the mean E* was shifted from ~0.61 to ~0.51, indicative of a closed-TL state with a probe-probe mean distance, R, of ~53 Å (Figs. 1d, bottom right). There was no indication of a higher-FRET, open-TL state within the temporal resolution of the experiment (~20 ms) in these experiments. Qualitatively identical results were obtained in experiments using each of two labelling sites in DNA (template-strand positions +10 and +12; Figs. 1d and S3). We conclude that, in a post-translocated TEC, in the presence of a saturating concentration of the complementary NTP, the TL is predominantly, potentially exclusively, in the closed-TL state. We conclude further that our smFRET assay enables unambiguous detection and differentiation of open-TL and closed-TL states in solution.

For TECs in the pre-translocated state, we observed unimodal E* distributions with mean E* of 0.62, indicative of an open-TL state (Fig. S4; Table S1), exactly as for TECs in the post-translocated state in the absence of an NTP (Fig. 1D, top right). There was no indication of a lower-FRET, closed-TL state within our temporal resolution (~20 ms). We conclude that, in a pre-translocated TEC state, the TL is predominantly, potentially exclusively, in an open-TL state.

### TL closing and opening occur on millisecond timescales

In a next set of experiments, we examined TL conformation in the presence of sub-saturating concentrations of the complementary NTP (ATP, using a template directing binding of ATP; Figs. 2 and S5). Upon addition of a sub-saturating concentration (20 μM) of the complementary NTP to post-translocated TECs, E* time trajectories for a subpopulation of molecules (~25%) showed transitions between two FRET states, indicating cycling between two TL conformational states (Fig. 2a). Hidden-Markov-Modelling (HMM) analysis of individual E* time trajectories identified two FRET states, one with E* of 0.60 (68%), corresponding to the open-TL state, and the other with E* of 0.46 (32%), corresponding to the closed-TL state (Fig. 2a, right; Table S1). From single-exponential fits of dwell-time distributions for the two FRET states, we estimated lifetimes of the TL-open state (900 ms) and TL-closed state (500 ms), and estimated rates of TL opening (k_open_) and TL closing (k_close_) (Figs. 2b-d and S5b-c). We next performed analogous experiments and analogous dwell-time analyses over a range of sub-saturating concentrations of complementary NTP (10 μM, 20 μM, 40 μM, and 80 μM complementary NTP; Figs. 2b-d; S5). The results revealed that, with increasing concentrations of complementary NTP, k_close_ increases and k_open_ remains constant (Figs. 2c-d and S5). We conclude that binding of a complementary NTP induces TL closure, that TL-closing events correspond to NTP-binding events, and that TL-opening events correspond to NTP unbinding events. From the ATP-concentration-dependence of k_close_, we estimate the on-rate for ATP, k_on,ATP_, as 0.045 μM^−1^s^−1^; from the mean value of k_open_ and the assumption that TL opening is fast relative to ATP dissociation, we estimate the off-rate for ATP, k_off,ATP_, as 2 s^−1^; and, from the association and dissociation rate for ATP, we estimate the equilibrium dissociation constant for ATP, K_d,ATP_, as 45 μM (Fig. 2d). The estimated value of K_d,ATP_ (45 μM) is in excellent agreement with the published value of K_d,ATP_ (44.6 μM^−1^s^−1^) (*46*).

**Fig. 2.**
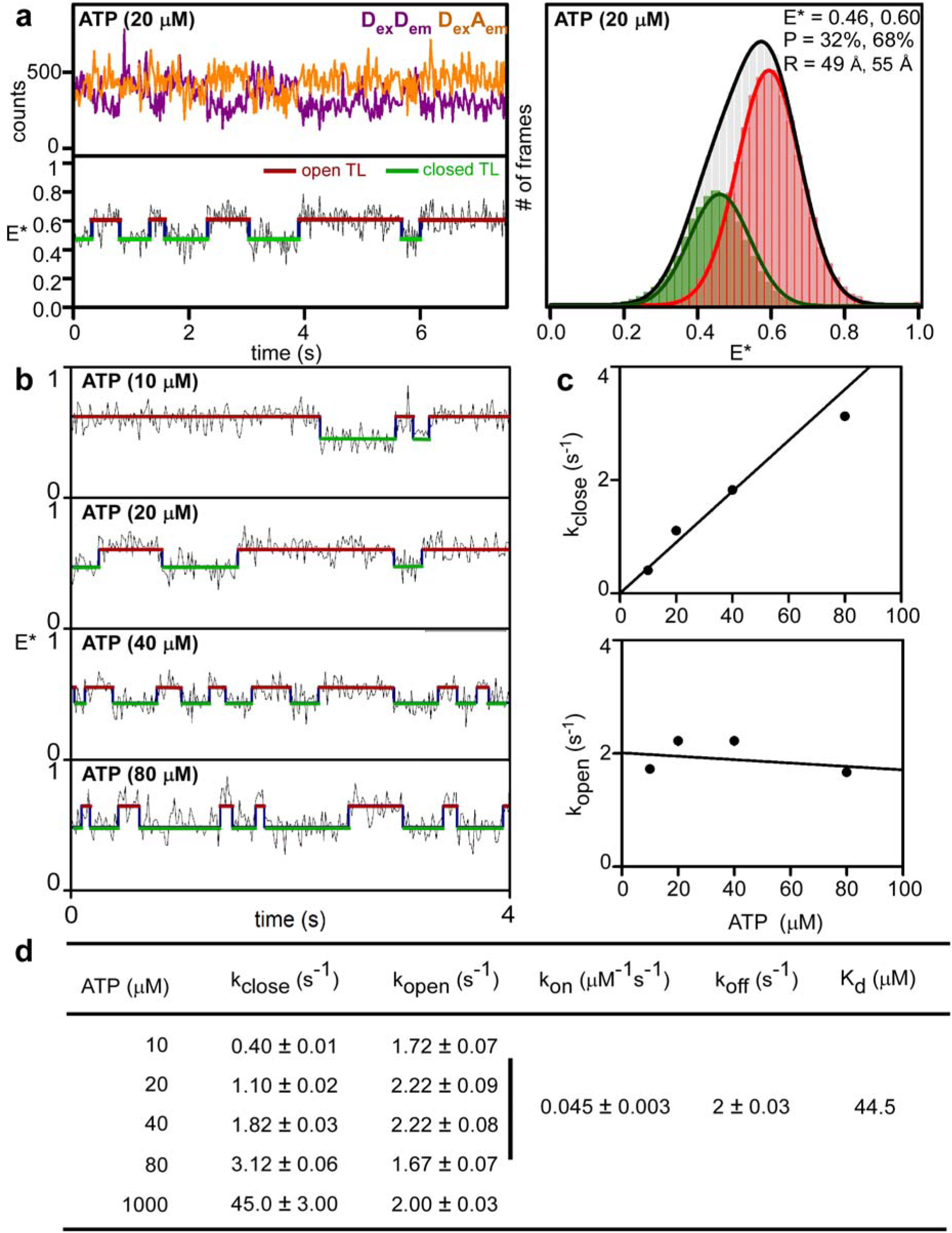
TL closing and opening occur on millisecond timescales. **a,** smFRET data for TEC in post-translocated state in presence of sub-saturating concentration of complementary NTP (20 μM ATP). Left top, representative time trace of donor emission (purple) and acceptor emission (orange). Left, bottom, representative time trace of donor-acceptor FRET efficiency, E*, showing hidden-Markov-model (HMM)-assigned open-TL states (red), closed-TL states (green), and interstate transitions (blue). Right, histograms and Gaussian fits of E*. P, subpopulation percentage; R, mean donor-acceptor distance. **b,** smFRET data for TEC in post-translocated state in presence of each of four sub-saturating concentrations of complementary NTP (10, 20, 40, and 80 μM ATP). Colors as in left bottom subpanel of a. **c,** ATP-concentration dependences of TL-closing rate (k_close_; top) and TL-opening rate (k_open_; bottom). **d,** TL-closing rate (k_close_), TL-opening rate (k_open_), ATP on-rate (k_on_), ATP off-rate (k_off_), and ATP equilibrium dissociation constant (K_d_) from experiments of a-c.

### TL closing and opening can provide a checkpoint for NTP complementarity, provide a checkpoint for NTP identity, and serve as a target for inhibitors

In a next set of experiments, we assessed effects of non-complementary NTPs on TL conformation (GTP, UTP, and CTP in experiments using a template directing binding of ATP; Fig. 3a; Table S1). In presence of saturating amounts of a non-complementary NTP, we observed unimodal E* distributions with E* of ~0.60, corresponding to the open-TL state (Fig. 3a; Table S1). There was no indication of a lower-FRET, closed-TL state within the temporal resolution of the experiment (~20 ms) in presence of any noncomplementary NTP (Fig. 3a). Analogous results were obtained in analogous experiments using templates directing binding of GTP (ATP, UTP, and CTP as non-complementary NTPs), directing binding of UTP (ATP, GTP, and CTP as non-complementary NTPs), and directing binding of CTP (ATP, GTP, and UTP as non-complementary NTPs) (Fig. S6). We conclude that the TL closes only with a complementary NTP and thus that TL closing can serve as a checkpoint for NTP complementarity.

**Fig. 3.**
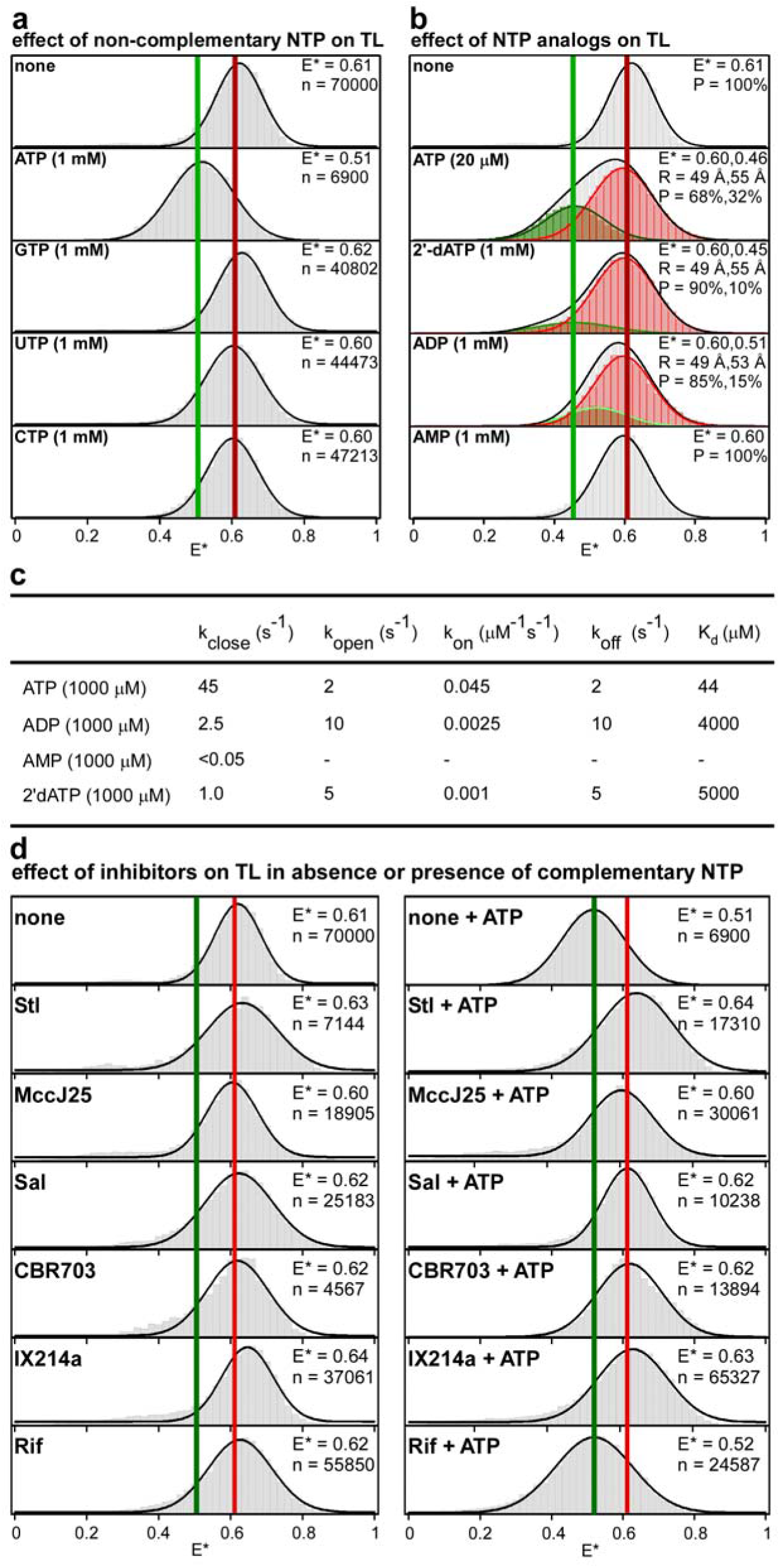
TL closing and opening can provide a checkpoint for NTP complementarity, provide a checkpoint for NTP identity, and serve as a target for inhibitors. **a,** Effects of non-complementary NTPs (GTP, UTP, and CTP for template directing incorporation of A) on TL conformation. Colors as in right subpanel of Fig. 1d. **b,** Effects of NTP analogues (dATP, ADP, and AMP) on TL conformation. Colors as in right subpanel of Fig. 2a. **c,** TL-closing rate (k_close_), TL-opening rate (k_open_), nucleotide association rate (k_on_), nucleotide dissociation rate (k_off_), and nucleotide equilibrium dissociation constant K_d_ from experiments of Fig. 3b. **d,** Effects of small-molecule inhibitors of RNAP on TL conformation in absence of NTP (left subpanels) and in presence of saturating concentration of complementary NTP (1 mM ATP; right subpanels) Stl, streptolydigin; MccJ25, microcin J25; Sal, salinamide A; CBR 703, CBR hydroxamidine CBR703; IX214a, Nα-aroyl-N-aryl-phenylalaninamide IX-214a; Rif, rifampin. Colors as in right subpanel of Fig. 1d.

We next assessed effects of NTP ribo/deoxyribo identity on TL conformation (ATP vs. dATP, using a template directing binding of ATP; Figs. 3b-c). The addition of dATP to 1 mM elicited only a very small change in the E* distribution, indicating that dATP is much less effective than ATP in inducing TL closure (Fig. 3b). Dwell-time analyses analogous to those performed with ATP (see previous section) indicated that the equilibrium dissociation constant for dATP is ~100 times higher than the equilibrium dissociation constant for ATP (5,000 μM vs 45 μM; Figs. 1D and 3C). We conclude that dATP induces TL closure only ~1/100 as potently as ATP and thus that TL closure can serve as a checkpoint for NTP ribo/deoxyribo identity.

We next assessed effects of NTP tri/di/monophosphate identity on TL conformation (ATP vs. ADP vs. AMP, using a template directing binding of ATP; Fig. 3b-c). The addition of ADP to 1 mM elicited only a very small change in the E* distribution, and the addition of AMP to 1 mM elicited no change in the E* distribution, indicating that ADP is much less effective than ATP in inducing TL closure and that AMP is ineffective in inducing TL closure (Fig. 3b). HMM analysis of individual E* time trajectories for experiments with 1 mM ADP identified two FRET states: one with E* of 0.60 (85%), corresponding to the open-TL state, and the other with E* of 0.51 (15%), corresponding to a partially-closed-TL state. The observation of a partially-closed-TL state in the presence of ADP suggests that ADP may bind to the RNAP active center and induce or stabilize an intermediate TL conformation with probe-probe distance of ~53 Å (Figs. 3b; Table S1). Dwell-time analyses for the open-TL and partially-closed-TL states analogous to those performed with ATP (see previous section) indicated that the equilibrium dissociation constant for ADP is ~100 times higher than the equilibrium dissociation constant for ATP (4,000 μM vs ~45 μM; Figs. 1d and 3c). We conclude that ADP induces TL closure only ~1/100 as potently as ATP, that AMP does not induce TL closure, and thus that TL closure can serve as a checkpoint for NTP tri/di/monophosphate identity.

We next evaluated effects of five small-molecule RNAP inhibitors that, based on crystal structures, interact with sites on RNAP that include or overlap the TL (Fig. 3d). Streptolydigin (Stl) interacts with the RNAP bridge helix C-terminal hinge and the TL, making direct contacts with the open-TL state that potentially stabilize the open-TL state (*9, 29–30*). Microcin J25 (MccJ25) interacts with the RNAP bridge helix N- and C-terminal hinges and the TL, making direct contacts with the open-TL state that potentially stabilize the open-TL state (*36*). Salinamide A (Sal) interacts with the RNAP bridge-helix N-terminal hinge in a manner that potentially sterically precludes TL closure (*31*). The CBR hydrazide CBR703 and the Nα-aroyl-N-aryl-phenylalaninamide IX214a interact with the RNAP bridge-helix N-terminal cap in a manner that potentially sterically precludes TL closure (*32–35*). We observed that none of the five small-molecule RNAP inhibitors significantly affected E* distributions in the absence of the complementary NTP (ATP in these experiments, using a template directing binding of A: Fig. 3d, left), but that all five affected E* distributions in the presence of the complementary NTP, completely inhibiting TL closure in the presence of the complementary NTP (Fig. 3d, right). The inhibition of TL closure in the presence of the complementary NTP was observed only for inhibitors that interact with sites on RNAP that include or overlap the TL; no such inhibition was observed for an inhibitor, rifampin (Rif), that interacts with a site on RNAP that does not include or overlap the TL (*47*; Fig. 3d). We conclude that the small-molecule inhibitors Stl, MccJ25, Sal, CBR703, and IX214a all inhibit TL closure in solution, and we conclude that the TL is a functional target for at least five classes of inhibitors in solution. We note that the smFRET assay of this report potentially could be adapted for high-throughput screening of TL-targeting small-molecule RNAP inhibitors.

### One TL closing-opening cycle typically occurs for each nucleotide addition in transcription elongation

We next monitored TL conformational cycling during real-time, active transcription elongation. To monitor TL conformational cycling during real-time, active transcription elongation, we prepared and analyzed TECs having a first fluorescent probe incorporated at a site in the RNAP TL (β’ residue 942; same site as in preceding sections) and having a second fluorescent probe incorporated at a reference site in RNAP (β residue 267; reference site selected to result in a large difference in probe-probe distance, and a corresponding large predicted difference in smFRET, for the open-TL state vs. the closed-TL state; Figs. 4a and S8a). We used a procedure comprising: (i) incorporation of two fluorescent probes, DyLight 550 and Dylight 650, one serving as donor and the other as acceptor, at RNAP β’ residue 942 and RNAP β residue 267, by use of unnatural-amino-acid mutagenesis and Staudinger ligation (*37, 39–40*); (ii) assembly of a doubly labelled TEC from the resulting doubly labelled RNAP and a nucleic-acid scaffold comprising a template-strand DNA oligonucleotide programming one, two, three, or four additions of A, a nontemplate-strand DNA oligonucleotide, and an RNA oligonucleotide; (iii) immobilization of the doubly labelled TEC, through a hexahistidine tag on RNAP, on an anti-hexahistidine tag-antibody-functionalized surface; and (iv) measurement of smFRET using TIRF-ALEX (Figs. 4a and S7).

To validate the use of this new class of constructs, we performed smFRET experiments analogous to those in Figs. 1–2, assessing a construct of this class that directed binding of ATP and that contained a non-extendable 3’-deoxyribonucleotide-containing RNA (Figs. S1c and S8). In the absence of a complementary NTP, we observed a unimodal E* distribution with mean E* of 0.59, indicative of an open-TL state with a probe-probe mean distance, R, of 53 Å (Fig. S8a, top; Table S2), and, in the presence of a saturating concentration of the complementary NTP (1 mM ATP), we observed a unimodal E* distribution with mean E* of 0.48, indicative of a closed-TL state with a probe-probe mean distance, R, of 58 Å (Fig. S8b, bottom; Table S2). In the presence of sub-saturating concentrations of the complementary NTP (10 μM, 20 μM, 40 μM, and 80 μM ATP), we observed transitions between two FRET states: one with E* of 0.59, corresponding to the open-TL state, and the other with E* of 0.48, corresponding to the closed-TL state (Fig. S8c; Table S2). Dwell-time analyses, performed as in the preceding sections, yielded estimates of k_on,ATP_ (0.038 μM^−1^s^−1^), k_off,ATP_ (biexponential fit; 17 s^−1^ and 2.4 s^−1^), and K_d,ATP_ (biexponential fit; 450 μM and 63 μM; Fig. S8C-D). The results demonstrate that this new class of constructs enables detection and differentiation of TL-open and TL-closed states in solution. We next assessed active transcription elongation, assessing constructs of this class that directed one addition of A, initiating transcription elongation by addition of ATP to 5 μM, and monitoring smFRET for 15 s (Figs. 4b, S1c, and S9; observation time limited to ~15 s by probe photobleaching at longer times). Approximately 25% of the molecules showed an unambiguous TL closing/opening cycle (characterized by a decrease in E* from ~0.6 to ~0.5 followed by an increase in E* from ~0.5 to ~0.6, each showing anti-correlated changes in signals in donor and acceptor channels) within the 15 s observation time (Fig. 4b). We extracted dwell-time distributions for the two FRET states, estimated lifetimes of the TL-open state (1,030 ms) and the TL-closed state (20 ms, 97% and 238 ms, 3%; biexponential fit), and estimated rates of TL opening (k_open_) and TL closing (k_close_) (Fig. S9A).

**Fig. 4.**
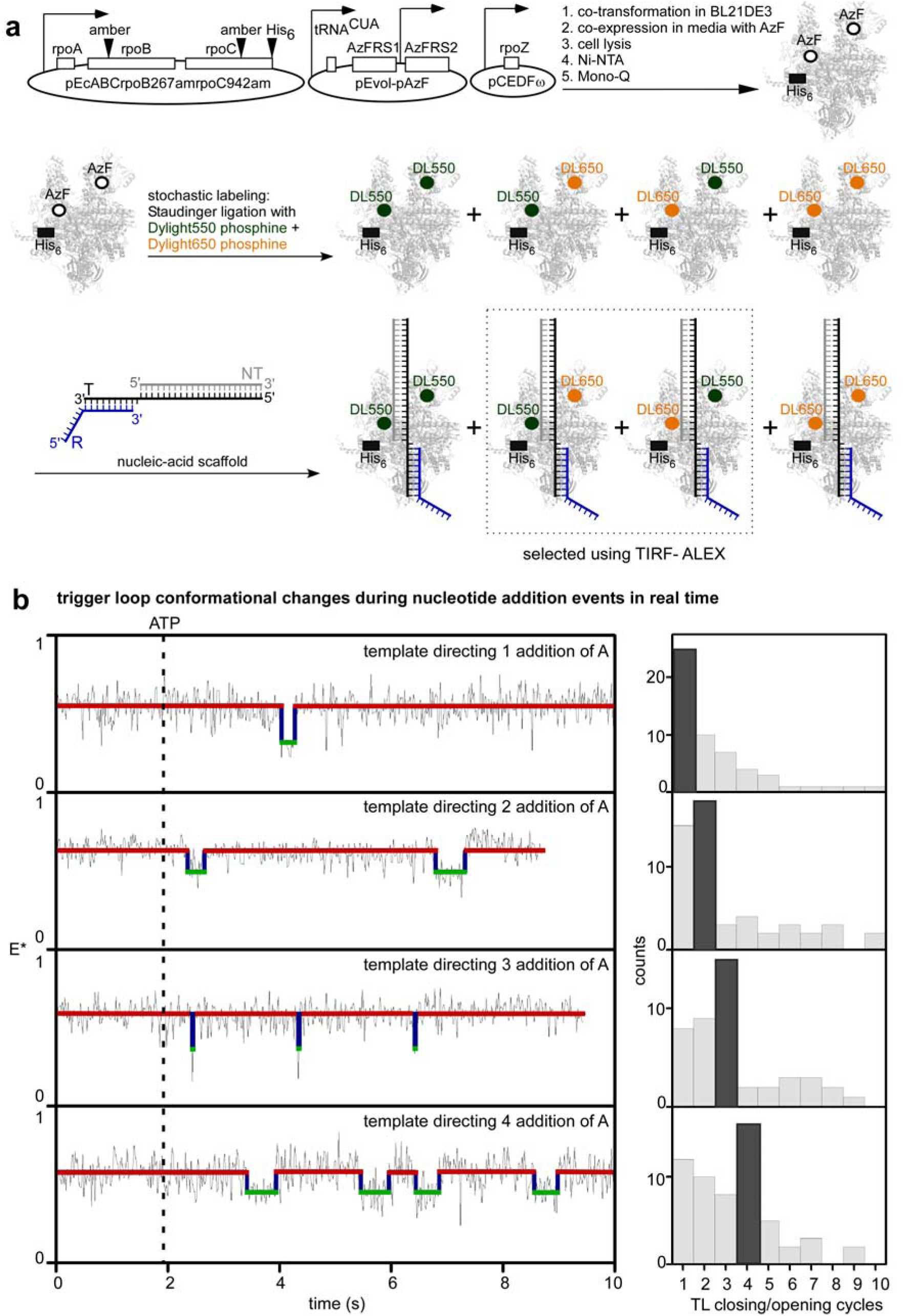
One TL closing-opening cycle typically occurs in each nucleotide addition in transcription elongation. **a,** Use of unnatural-amino-acid mutagenesis (first reaction arrow), Staudinger ligation with Dylight 550-phosphine and Dylight 650-phosphine (second reaction arrow), TEC reconstitution (third reaction arrow), and total-internal-reflection fluorescence microscopy with alternating-laser excitation microscopy (TIRF-ALEX) to measure smFRET between first fluorescent probe at tip of RNAP TL and second fluorescent probe at reference site elsewhere in RNAP (see Methods). Green filled circles, fluorescent probe Dylight 550; orange filled circles, fluorescent probe Dylight 650. Other colors as in Fig. 1c. **b,** smFRET data for TEC engaged in active real-time transcription elongation on templates directing addition of 1, 2, 3, or 4 additions of A. Left subpanels, representative time traces of donor-acceptor FRET, E*. Black dashed line, addition of ATP. Other colors as in left bottom subpanel of Fig. 2a. Right subpanels, histograms showing numbers of detected TL closing/opening cycles. Note one-for-one correlation between the modal number of TL closing/opening events detected for a template (dark gray bar for each template) and the number of nucleotide additions directed by the template.

We note that the mean dwell time for the TL-closed state in experiments assessing active transcription (1/k_open_; 20 ms^;^ Fig. S9a) is only ~1/3 the mean dwell time for the TL-closed state in experiments assessing NTP binding (1/ k_open_; ~55 ms; Fig. S8d). We suggest that this difference relates to the fact that the mean dwell time for the TL-closed state in experiments assessing active transcription (Fig. 4) corresponds to reactions from TL closing through *either* phosphodiester-bond formation and TL opening, *or* NTP dissociation and TL opening, *whichever comes first*, whereas the mean dwell time for the TL-closed state in experiments assessing NTP binding (Fig. 2) corresponds to reactions from TL closing through NTP dissociation and TL opening. Assuming that the mean dwell time for the TL-closed state in experiments assessing active transcription at 5 μM ATP approximates the mean time for reactions from TL closing through phosphodiester-bond formation and TL opening (1/k_open_; 20 ms; Fig. S9a), and, assuming that the mean dwell time for the TL-open state in experiments assessing active transcription approximates at 5 μM ATP approximates the mean time for the other reactions of the nucleotide-addition cycle (1/k_close_; 1,030 ms; Fig. S9a), we estimate that the mean total duration of a nucleotide-addition cycle at 5 μM ATP is 1,050 ms. This estimate is in excellent agreement with the reported value for the mean total duration of a nucleotide-addition cycle at 5 μM ATP: ~1,000 ms (*4*).

We next *counted* the number of TL closing/opening events for constructs of this class that directed one, two, three, or four additions of A, initiating transcription elongation by addition of ATP to 5 μM at t = 0, and monitoring smFRET for 15 s (Figs. 4b, S1c, and S9b). We observed a remarkable, one-for-one correlation between the modal number of TL closing/opening events detected for a nucleic-acid scaffold and the number of nucleotide additions directed by the nucleic-acid scaffold (Fig. 4b); the modal numbers of TL closing/opening events detected were one, two, three, and four for nucleic-acid scaffolds directing, respectively, one, two, three, and four nucleotide additions (Fig. 4b, right). We conclude that one cycle of TL closing/opening typically occurs for each nucleotide-addition step in transcription elongation.

Most, possibly all, cases in which the number of detected TL closing/opening events was lower than the modal number (~30% of molecules) may represent cases where events occurred but were unresolved within our ~20 ms temporal resolution; analysis of dwell-time distributions indicates that ~60% of events are likely to have durations below 20 ms (Fig. S9a). At least some cases in which the numbers of detected TL closing/opening events was higher than the modal number (~30% of molecules) may represent cases where events occurred that did not result in net nucleotide addition, such as, for example, events where nucleotide addition was followed by nucleotide removal by pyrophosphorolysis or hydrolysis (*2*). Accordingly, we suggest that one cycle of TL closing/opening may *always, or almost always,* occur for each net nucleotide addition in transcription elongation.

### Prospect

Our results demonstrate the direct detection of TL conformation and conformational cycling in solution (Figs. 1–2); show that the TL is open in the absence of an NTP in solution (Figs. 1d and 2); show that the TL closes upon binding an NTP in solution (Figs. 1d and 2); show that TL closing and opening occur on the millisecond time scale in solution (Fig. 2); show that the TL serves as a checkpoint for NTP complementarity, NTP ribo/deoxyribo identity, and tri/di/monophosphate identity (Fig. 3a-c); show that the TL serves as a target for five classes of small-molecule RNAP inhibitors (Fig. 3d); and, most important, shows that, in most, and possibly all, cases, one TL closing/opening cycle occurs in each nucleotide-addition step during transcription elongation (Fig. 4). Our results provide direct support for the hypothesis from crystal structures (>*2, 5, 8–11*) that the RNAP TL undergoes two large-amplitude conformational changes--first closing, with movement by ~15-25 Å, and then opening, with movement by ~15-25 Å--in each millisecond-timescale nucleotide-addition step in transcription elongation. Our results also provide direct support for the hypothesis from crystal structures (*2, 5, 8–11*) that RNAP TL conformational cycling is functionally important for substrate specificity and catalysis. Our results imply--in view of the TL’s location near the center of the RNAP molecule, its conformational cycling, and its functional importance--that the TL is the veritable “beating heart” of RNAP.

Because the TL is conserved in RNAP from all living organisms, our conclusions regarding TL conformational cycling and TL functional importance are likely to be valid for RNAP from bacteria through humans, and our smFRET approach for analysis of TL conformation is likely to be applicable to RNAP from bacteria through humans.

We note that combining our smFRET approach with optical-tweezers or nanopore-tweezers approaches able to detect “stepping” of RNAP at single-base-pair resolution (*4, 48*) could enable analysis of TL conformation and TL conformational cycling as a function of template location and template sequence in transcription elongation, transcriptional pausing, and transcription termination.

## Acknowledgements

R.H.E. was supported by NIH grant GM041376. A.N.K. was supported by Wellcome Trust grant 110164/Z/15/Z, and the UK Biotechnology and Biological Sciences Research Council grants BB/H01795X/1 and BB/J00054X/1. We thank W. Fenical, and E. Steinbrecher for compounds.

## Author Contributions

R.H.E conceived the project. A.M., A.N.K., and R.H.E. designed experiments. A.M. and M.L. prepared RNAP derivatives. A.M. performed single molecule experiments. A.M., R.H.E, and A.N.K analyzed data. A.M, A.N.K., and R.H.E. wrote the paper.

## Supplement: Materials and Methods

### RNAP derivatives

For experiments in Figs. 1–3 and S2–S6, fluorescent-probe-labelled, hexahistidine-tagged *Escherichia coli* RNAP core enzyme was prepared using unnatural-amino-acid mutagenesis (*49*) of co-expressed genes encoding RNAP β’, β, α, and ω subunits to afford an RNAP core enzyme derivative containing 4-azido-L-phenylalanine (AzF) at position 942 of β’, followed by azide-specific Staudinger ligation to incorporate the fluorescent probe Dylight 550 (DL550) at position 942 of β’, as follows (Figs. 1c, S2; *50-52*): Single colonies of *E. coli* strain BL21(DE3) (Millipore) co-transformed with plasmid pEcABC-rpoC942am-His_6_ [constructed from plasmid pEcABC-His_6_ (*53*) by use of site-directed mutagenesis (QuikChange Site-Directed Mutagenesis Kit; Agilent) to replace *rpoC* codon 942 by an amber codon], plasmid pCDFω (*54*), and plasmid pEVOL-pAzF (*49*) were used to inoculate 20 ml LB broth (*55*) containing 100 μg/ml ampicillin, 50 μg/ml kanamycin, and 35 μg/ml chloramphenicol, and cultures were incubated 16 h at 37°C with shaking. Culture aliquots (2×10 ml) were used to inoculate LB broth (2×1 L) containing 2 mM AzF (Chem-Impex International), 100 μg/ml ampicillin, 50 μg/ml kanamycin, and 35 μg/ml chloramphenicol; cultures were incubated at 37°C with shaking until OD_600_ = 0.6; L-arabinose was added to 0.2% and IPTG was added to 1 mM; and cultures were further incubated 16 h at 16°C with shaking. Cells were harvested by centrifugation (4,000 × g; 20 min at 4°C), re-suspended in 20 ml buffer A (10 mM Tris-HCl, pH 7.9, 200 mM NaCl, and 5% glycerol), and lysed using an EmulsiFlex-C5 cell disrupter (Avestin). The lysate was cleared by centrifugation (20,000 × g; 30 min at 4°C), precipitated with polyethyleneimine (Sigma-Aldrich) as in (*56*), and precipitated with ammonium sulfate as in (*56*). The precipitate was dissolved in 30 ml buffer A and loaded onto a 5 ml column of Ni-NTA-agarose (Qiagen) pre-equilibrated in buffer A, and the column was washed with 50 ml buffer A containing 10 mM imidazole and eluted with 25 ml buffer A containing 200 mM imidazole. The sample was further purified by anion-exchange chromatography on Mono Q 10/100 GL (GE Healthcare; 160 ml linear gradient of 300-500 mM NaCl in 10 mM Tris-HCl, pH 7.9, 0.1 mM EDTA, and 5% glycerol; flow rate = 2 ml/min). Fractions containing AzF-derivatized hexahistidine-tagged *E. coli* RNAP core enzyme were pooled, concentrated to ~1 mg/ml using 30 kDa MWCO Amicon Ultra-15 centrifugal ultrafilters (EMD Millipore), and stored in aliquots at −80°C. A reaction mixture containing 10 μM AzF-derivatized hexahistidine-tagged *E. coli* RNAP core enzyme and 250 μM DL550 phosphine (Thermo Fisher Scientific; Cat. no. 88910) in 1 ml buffer B (50 mM Tris-HCl, pH 7.9, 100 mM KCl, 5% glycerol, and 2% dimethylformamide) was incubated 1 h at 15°C, incubated 16 h on ice, and subjected to 5 cycles of buffer exchange (dilution with 5 ml buffer B, followed by concentration to 0.5 ml using 30 kDa MWCO Amicon Ultra-15 centrifugal ultrafilters). The sample was further purified by gel-filtration chromatography on HiLoad 16/60 Superdex 200 prep grade (GE Healthcare) pre-equilibrated in buffer C (20 mM Tris-HCl, pH 8.0, 100 mM NaCl, 5 mM MgCl_2_, 1 mM β-mercaptoethanol, and 5% glycerol) and eluted in buffer C. Fractions containing fluorescent-probe-labelled, hexahistidine-tagged *E. coli* RNAP core enzyme were pooled, concentrated to 1 mg/ml in buffer C using 30 kDa MWCO Amicon Ultra-15 centrifugal ultrafilters, and stored in aliquots at −80°C.

For experiments in Figs. 4 and S7–S9, fluorescent-probe-labelled, hexahistidine-tagged *E. coli* RNAP core enzyme was prepared using unnatural-amino-acid mutagenesis (*49*) of co-expressed genes encoding RNAP β’, β, α, and ω subunits to afford an RNAP core enzyme derivative containing AzF at position 942 of β’ and position 267 of β, followed by stochastic azide-specific Staudinger ligation to incorporate the fluorescent probes Dylight 550 (DL550) and Dylight 650 (DL650) at position 942 of β’ and position 267 of β (Figs. 4a, S7; *50-51*). The RNAP core enzyme derivative containing AzF at position 942 of β’ and position 267 of β was prepared as described in the preceding paragraph for the RNAP core derivative containing AzF at position 942 of β’, but using plasmid pEcABC-rpoB267am;rpoC942am-His_6_ [constructed from plasmid pEcABC-rpoC942am-His_6_ by use of site-directed mutagenesis (QuikChange Site-Directed Mutagenesis Kit; Agilent) to replace *rpoB* codon 267 by an amber codon] in place of plasmid pEcABC-rpoC942am-His_6._. A reaction mixture containing 10 μM AzF-derivatized, hexahistidine-tagged *E. coli* RNAP core enzyme, 1000 μM Dylight 550 phosphine (Thermo Fisher Scientific; Cat. no. 88910), and 500 μM Dylight 650 phosphine (Thermo Fisher Scientific; Cat. no. 88911) in 1 ml buffer B (50 mM Tris-HCl, pH 7.9, 100 mM KCl, 5% glycerol, and 2% dimethylformamide) was incubated 1 h at 15°C, incubated 16 h on ice, subjected to 5 cycles of buffer exchange (dilution with 5 ml buffer B, followed by concentration to 0.5 ml) using 30 kDa MWCO Amicon Ultra-15 centrifugal ultrafilters (EMD Millipore), and stored in aliquots at −80°C.

Efficiencies of incorporation of fluorescent probes were determined from UV/Vis-absorbance measurements and were calculated as::

> concentration of product = [A_280_ −ε_DL550,280_ (A_DL550,562_/ε_Cy3B,562_) − ε_DL650,280_ (A_DL650,660_/ε_DL650,660_)]/ε_P,280_
>
> DL550 labelling efficiency = 100% [(A_DL550,562_/ε_DL550,562_)/(concentration of product)]
>
> DL650 labelling efficiency = 100% [(A_DL650,660_/ε_DL650,660_)/(concentration of product)]

where A_280_ is the measured absorbance at 280 nm, A_DL550,562_ is the measured absorbance at the long-wavelength absorbance maximum of DL550 (562 nm), A_DL650,660_ is the measured absorbance at the long-wavelength absorbance maximum of DL650 (660 nm), ε_P,280_ is the molar extinction coefficient of RNAP core enzyme at 280 nm (198,500 M^−1^ cm^−1^), ε_DL550,280_ is the molar extinction coefficient of DL550 at 280 nm (12,090 M^−1^ cm^−1^), ε_DL650,280_ is the molar extinction coefficient of DL 650 at 280 nm (9,250 M^−1^ cm^−1^), ε_DL550,562_ is the extinction coefficient of DL550 at its long-wavelength absorbance maximum (150,000 M^−1^ cm^−1^), and ε_DL650,660_ is the extinction coefficient of DL650 at its long-wavelength absorbance maximum (250,000 M^−1^ cm^−1^). Labelling efficiencies were ~60% for DL550 for the singly labelled RNAP derivative (Figs. S2) and ~50% for DL550 and ~80% for DL650 for the doubly labelled RNAP derivative (Fig. S7).

Specificities of incorporation of fluorescent probes were determined from the observed labelling efficiencies of (i) the labelling reaction with the AzF-derivatized, hexahistidine-tagged *E. coli* RNAP core enzyme and (ii) a control labelling reaction with non-AzF-derivatized, hexahistidine-tagged *E. coli* RNAP core enzyme (prepared as described in *57*), and were calculated as:

labelling specificity =100% [1-[(labelling efficiency with P)/ (labelling efficiency with AzF-P)]

where AzF-P is AzF-derivatized, hexahistidine-tagged *E. coli* RNAP core enzyme, and P is non-AzF-derivatized, hexahistidine-tagged *E. coli* RNAP core enzyme. Labelling specificities were >90% (Figs. S2, S7).

Transcriptional activities of labelled RNAP derivatives were determined using fluorescence-detected transcription assays as described in (*58*).

**_σ_**70

*E. coli* σ^70^ was prepared as in (*57*).

### Nucleic acids

Oligodeoxyribonucleotides (Integrated DNA Technologies, Inc) and oligoribonucleotides (Trilink, Inc) were dissolved in nuclease-free water (Ambion, Inc) to a final concentration of 100 mM and stored at − 20°C. Oligodeoxyribonucleotides were labelled with Alexa Fluor 647 N-hydroxysuccinimide (NHS) ester (Molecular Probes) as described (*59*).

### Nucleic-acid scaffolds

Nucleic-acid scaffolds (Fig. S1) were prepared as follows: Nontemplate-strand oligodeoxyribonucleotide (3 mM), template-strand oligodeoxyribonucleotide (2 mM), and oligoribonucleotide (8 mM) in 50 μl 10 mM Tris-HCl, pH 7.9 and 0.2 M NaCl were heated 5 min at 95°C, cooled to 25°C in 2°C steps with 1 min per step using a thermal cycler (Applied Biosystems), and stored at −20°C.

### Small molecules

NTPs (Thermo Fisher Scientific, Cat. No. R0481), dATP (Thermo Fisher Scientific, Cat. No. R0141), ADP (Sigma-Aldrich), and AMP (Sigma-Aldrich) were diluted in nuclease-free water (Ambion, Inc) to final concentrations of 25 mM and stored in aliquots at −80°C.

Streptolydigin (Stl) was the kind gift of Dr. E. Steinbrecher (Upjohn-Pharmacia, Kalamazoo, MI), and Salinamide A (Sal A) was the kind gift of Dr. W. Fenical (The Scripps Research Institute, La Jolla, CA). Rif was purchased from Sigma-Aldrich, and CBR703 was purchased from Maybridge. MccJ25 was prepared as in (*60*), and IX214A was prepared as in (*61*).

### Transcription elongation complexes (TECs)

Fluorescent-probe-labelled, hexahistidine-tagged transcription elongation complexes (TECs) were prepared as follows: Reaction mixtures containing 10 nM single fluorescent-probe-labelled, hexahistidine-tagged *E. coli* RNAP core enzyme and 100 nM fluorescent-probe labelled nucleic-acid scaffold (for experiments in Figs.1c, 2, 3 and Figs. S3–S6; sequences in Figs. S1a-b), or 10 nM fluorescent-probe-doubly-labelled, hexahistidine-tagged *E. coli* RNAP core enzyme and 100 nM nucleic-acid scaffold (for experiments in Figs. 4 and Figs. S8–S9; sequences in Figs. S1c) in 0.5 ml KG7 (40 mM HEPES-NaOH, pH 7.0, 100 mM potassium glutamate, 10 mM MgCl_2_, 1 mM dithiothreitol, 100 μg/ml bovine serum albumin, and 5% glycerol) were incubated 15 min at 22°C. Reaction mixtures then were concentrated to 0.05 ml and subjected to by 3 cycles of buffer exchange (dilution with 0.5 ml buffer KG7, followed by concentration to 0.05 ml) using 100 kDa MWCO Amicon Ultra-0.5 centrifugal filters (Millipore). TECs prepared using this procedure were stable for up to 24 h on ice.

### smFRET using confocal-ALEX

Confocal-ALEX experiments were performed essentially as described (*50, 59*). A green laser (532 nm; Compass 215M-20; Coherent) was used for direct excitation of the donor, and a red laser (638 nm; Radius 635-25; Coherent,) was used for direct excitation of the acceptor (*50*). Lasers were operated at continuous-wave excitation intensities of 120 μW at 532 nm and 80 μW at 638 nm and were alternated at 25 μs intervals using an acousto-optical modulator (Neos Technologies, Inc.). Fiber-coupled collimated beams were directed to an Olympus IX71 inverted microscope (Olympus America, Inc.), reflected by a beam splitter, and focused into the sample through a 60x oil-immersion objective. Fluorescence emission from the sample was collected through the objective, filtered through a 100 μm pinhole, spectrally split by a dichroic mirror, and focussed onto two avalanche photodiode detectors (APD; SPCM-AQR-15; Perkin-Elmer).

For experiments in Figs. S3 and S6, TECs were diluted to a final concentration of 0.1 nM in 50 μl KG7-trolox [KG7 containing 2 mM Trolox (Sigma Aldrich)] containing the following nucleotides: (i) none; (ii) 1 mM ATP; (iii) 1 mM GTP; (iv) 1 mM CTP, or (v) 1 mM UTP. Following incubation 5 min at 22°C, smFRET data collection was performed. Data-acquisition times ranged from 20-30 minutes.

Photons detected at the donor-emission channel upon donor excitation (F_DD_), acceptor-emission channel upon donor excitation (F_DA_), and acceptor-emission channel upon acceptor excitation (F_AA_) were extracted based on photon arrival times. The stoichiometry parameter (S) was calculated for each above-threshold photon burst, as follows (*50, 59, 62*):

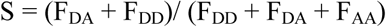

The donor-acceptor smFRET efficiency (E*) for each above-threshold, photon burst was calculated as follows (*50, 59, 62*):

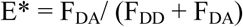

Two-dimensional E*-S plots were used to distinguish species containing donor only (D-only), acceptor only (A-only), and both donor and acceptor (D-A). For species containing both donor and acceptor (D-A), one-dimensional E* histograms were plotted and were fitted with Gaussian curves (Figs. S3 and S6). The resulting histograms provided equilibrium population distributions of E*.

### smFRET using TIRF-ALEX: sample preparation

Observation wells for smFRET experiments were prepared as described (*51–52*). Briefly, a biotin-PEG-passivated glass surface was prepared, functionalized with Neutravidin (Sigma Aldrich), and treated with biotinylated anti-hexahistidine monoclonal antibody (Penta-His Biotin Conjugate; Qiagen), yielding wells with (biotinylated anti-hexahistidine monoclonal antibody)-Neutravidin-biotin-PEG-functionalised glass floors. For experiments in Figs. 1c, 2–4 and Figs. S3–S6 and S8–9, fluorescent-probe-labelled, hexahistidine tagged TECs were immobilised in observation wells with (biotinylated anti-hexahistidine monoclonal antibody)-Neutravidin-biotin-PEG-functionalized glass floors, as follows: aliquots (30 μl) of 0.1 nM fluorescent-probe-labelled, hexahistidine tagged TEC in KG7 were added to the observation chamber and incubated 2-4 min at 22°C, solutions were removed, wells were washed with 2X30 μl KG7, and 30 μl imaging buffer [KG7 containing 2 mM Trolox (Sigma-Aldrich), 12.5 μM glucose oxidase (Sigma-Aldrich), 16 nM catalase (from bovine liver C30; Sigma-Aldrich), and 8 mM D-glucose] at 22°C was added. Immobilization densities typically were ~20-30 molecules per 10 μm × 12 μm field of view (assessed by manual counting). Immobilization specificities typically were >98% (assessed in control experiments omitting biotinylated anti-hexahistidine monoclonal antibody).

For experiments in Fig. 3d in the absence of NTPs, observation chambers containing immobilized fluorescent-probe-labelled, hexahistidine-tagged TECs were prepared as described above, except that (i) 1 nM fluorescent-probe-labelled, hexahistidine-tagged TEC was pre-incubated 5 min at 22°C in KG7 containing the specified concentration of inhibitor (none, 20 μM Rif, 50 μM Stl, 100 μM MccJ25, 20 μM Sal, 50 μM CBR703, or 200 μM IX214A) and then diluted 1:10 with KG7 containing the specified concentration of inhibitor before addition of aliquots to observation chambers, and (ii) the KG7 wash stocks contained the specified concentration of inhibitor, and (iii) the imaging buffer stocks contained the specified concentration of inhibitors.

For experiments in Figs. 1c, 2, 3a-c, S5, and S8 in the presence of NTPs or NTP analogs, observation chambers containing immobilized fluorescent-probe-labelled, hexahistidine-tagged TECs (prepared as described above) were supplemented to the specified final concentrations with 30 μl solutions of NTPs or NTP analogs in imaging buffer, reaction mixtures were incubated 3 min at 22°C, and data were collected.

For experiments in Figs. 4 and S9, observation chambers containing immobilized fluorescent-probe-labelled, hexahistidine-tagged TECs were prepared as described above with 30 μl imaging buffer in the observation chambers, data acquisition was started, and observation chambers were supplemented with 10μl of 20μM ATP in imaging buffer (yielding a final ATP concentration of 5 μM).

### smFRET using TIRF-ALEX: data collection and data analysis

smFRET experiments were performed using a custom-built objective-type total-internal-reflection fluorescence (TIRF) microscope (*63*). Light from a green laser (532 nm; Samba; Cobolt) and a red laser (635 nm; CUBE 635-30E, Coherent) was combined using a dichroic mirror coupled into a fiber-optic cable focused onto the rear focal plane of a 100x oil-immersion objective (numerical aperture 1.4; Olympus) and was displaced off the optical axis, such that the incident angle at the oil-glass interface of a stage-mounted observation chamber exceeded the critical angle, thereby creating an exponentially decaying evanescent wave (*64*). Alternating-laser excitation (ALEX; *59*) was implemented by directly modulating the green and red lasers using an acousto-optical modulator (1205C, Isomet).

Fluorescence emission was collected from the objective, was separated from excitation light using a dichroic mirror (545 nm/650 nm, Semrock) and emission filters (545 nm LP, Chroma; and 633/25 nm notch filter, Semrock), was focused on a slit to crop the image, and then was spectrally separated (using a dichroic mirror; 630 nm DLRP, Omega) into donor and emission channels focused side-by-side onto an electron-multiplying charge-coupled device camera (EMCCD; iXon 897; Andor Technology). A motorized x/y-scanning stage with continuous reflective-interface feedback focus (MS-2000; ASI) was used to control the sample position relative to the objective.

All data acquisition was carried out at 22°C. For all TIRF-ALEX experiments except those in Fig. 3D, laser powers were 4 mW (532 nm laser) and 0.75 mW (635 nm laser), and data were collected for 20 s using a frame rate of 1 frame per 20 ms. For experiments in Fig. 3D, laser powers were 1 mW (532 nm laser) and 0.3 mW (635 nm laser), and data were collected for 50 s using a frame rate of 1 frame per 100 ms.

Fluorescence-emission intensities in donor-emission (green) and acceptor-emission (red) channels were detected using the peak-finding algorithm of the MATLAB (MathWorks) software package Twotone-ALEX, as described (*63*). Peaks detected in both emission channels (i.e., peaks for molecules containing both donor and acceptor probes) were fitted with two-dimensional Gaussian functions to extract background-corrected intensity-vs.-time trajectories for donor-emission intensity upon donor excitation (I_DD_), acceptor-emission intensity upon donor excitation (I_DA_), and acceptor-emission intensity upon acceptor excitation (I_AA_), as described (*63*). Intensity-vs.-time trajectories were curated to exclude trajectories exhibiting I_DD_ <100 or >1,000 counts or I_AA_ <200 or >1,000 counts, trajectories exhibiting multiple-step donor or acceptor photobleaching, trajectories exhibiting donor or acceptor photobleaching in frames 1-20, trajectories exhibiting donor or acceptor photoblinking, trajectories exhibiting E* values < 0.3 (inferred to be donor-only complexes or improperly assembled complexes), and portions of trajectories following donor or acceptor photobleaching.

Intensity-vs.-time trajectories were used to calculate trajectories of apparent donor-acceptor smFRET efficiency (E*) and donor-acceptor stoichiometry (S), as described (*59, 62*):

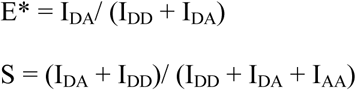

E*-vs.-S plots were prepared, S values were used to distinguish species containing only donor, only acceptor, and both donor and acceptor, and E* histograms were prepared for species containing both donor and acceptor, as described (*59, 62*), and fitted to Gaussian distributions in Origin (Origin Lab). The resulting histograms provide equilibrium population distributions of E* states and, for each E* state, define mean E* (Figs. 1C, 3a, S4, S5 and S8b; gray bars and inset).

E*-vs-time trajectories that, on visual inspection, exhibited transitions between distinct E* states and exhibited anti-correlated changes in DD (donor excitation-donor emission) and DA (donor excitation-acceptor emission) channels were identified. For experiments in Figs. 1C, S4 and S8, few traces (<3%) showed dynamic behavior. For experiments in Figs. 2, S5 and S8, ~10% to ~60% of traces showed dynamic behavior. Dynamic E*-vs-time trajectories were analyzed globally to identify E* states by use of Hidden Markov Modelling (HMM) as implemented the Matlab (MathWorks) software package ebFRET (*65*), essentially as described (*51, 65*). E*-vs-time trajectories were fitted to a two-state HMM model, E*-values from the fitted model were extracted, were plotted using Origin (Origin Lab), and were fitted to Gaussian distributions using Origin (Figs. 2b, 3b and S5; colored curves). The resulting histograms provide equilibrium population distributions of E* states and, for each E* state, define mean E* (Figs. 2b, 3b and S5; colored bars and inset).

E* values were corrected, and accurate donor-acceptor efficiencies (E_a_) and donor-acceptor distances (R) were calculated (Table S1) as described previously (*52*). From the accurate FRET efficiencies (E_a_), distances (R; Table S1) were estimated using:

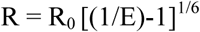

where R_0_ is the Főrster parameter for the donor/acceptor FRET pair [52 Å for DL550-DL650 FRET pair (*66*); 50 Å for DL550-Alexa647 FRET pair; calculated as: R_0_= 9780(n^−4^κ^2^Q_D_J)^1/6^ Å, where n is the refractive index of the medium, κ^2^ is the orientation factor relating donor and acceptor transition dipoles (approximated as 2/3, noting that all mean E values are <0.5; *67*), Q_D_ is the donor quantum yield, and J is the overlap integral of donor emission and acceptor excitation].

Dwell times for E* states were extracted from HMM fits to E*-vs-time trajectories and were binned and plotted as distribution histograms in Origin (Fig. S5). For experiments in Figs. 2, 3b and S5, the rate of TL closing (k_close_) and the rate of TL opening (k_open_) were estimated from single-exponential fits to the open-TL and closed-TL dwell-time-distribution histograms, respectively (Figs. 2d, 3c, and S5).

For experiments in Figs. 4, S8, and S9, the open-TL dwell-time-distribution histograms were fit to a single-exponential function, and the closed dwell-time-distribution histograms were fit to a bi-exponential function with most events corresponding to short dwells (~60 ms; ~90%) and some events corresponding to longer dwells (~400 ms; ~10%). The rate of TL opening (k_open_) and the rate of TL closing (k_close_) were estimated from exponential fits to closed and open dwell time distribution histograms, respectively, as described above (Fig. S8d and S9).

The on-rate for ATP binding, the off-rate for ATP unbinding and the equilibrium dissociation constant for ATP binding were estimated from the TL-closing and TL-opening rates (assuming TL closing events correspond to NTP-binding events, and TL-opening events correspond to NTP-unbinding events), as follows (Fig. 2e, 3c, S8d):

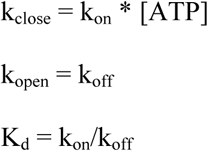

For experiments in Figs. 4 and S9, ~25% of molecules exhibited transitions between distinct E* states and exhibited anti-correlated changes in the DD (donor excitation-donor emission) and DA (donor excitation-acceptor emission) channels, upon addition of 5 μM ATP. One TL-closing step (transition from open-TL state to closed-TL state) through one TL-opening step (transition from closed-TL to open-TL state) was defined to constitute a TL closing-opening event. For each experiment in Fig. 4 (template directing addition of 1A, 2A, 3A, or 4A), a manual counting of numbers of TL closing-opening events was performed, and numbers of TL closing-opening events were plotted as a histogram in Origin. Rare complexes (<1%) that showed TL closing-opening events prior to ATP addition were excluded from the analysis. TL-closing and TL-opening rates were estimated as described above (Fig. S9). Molecules which did not show any transitions may include time traces where all transitions are missed or molecules which were unresponsive to addition of 5 μM ATP.

---

**Fig. S1.**
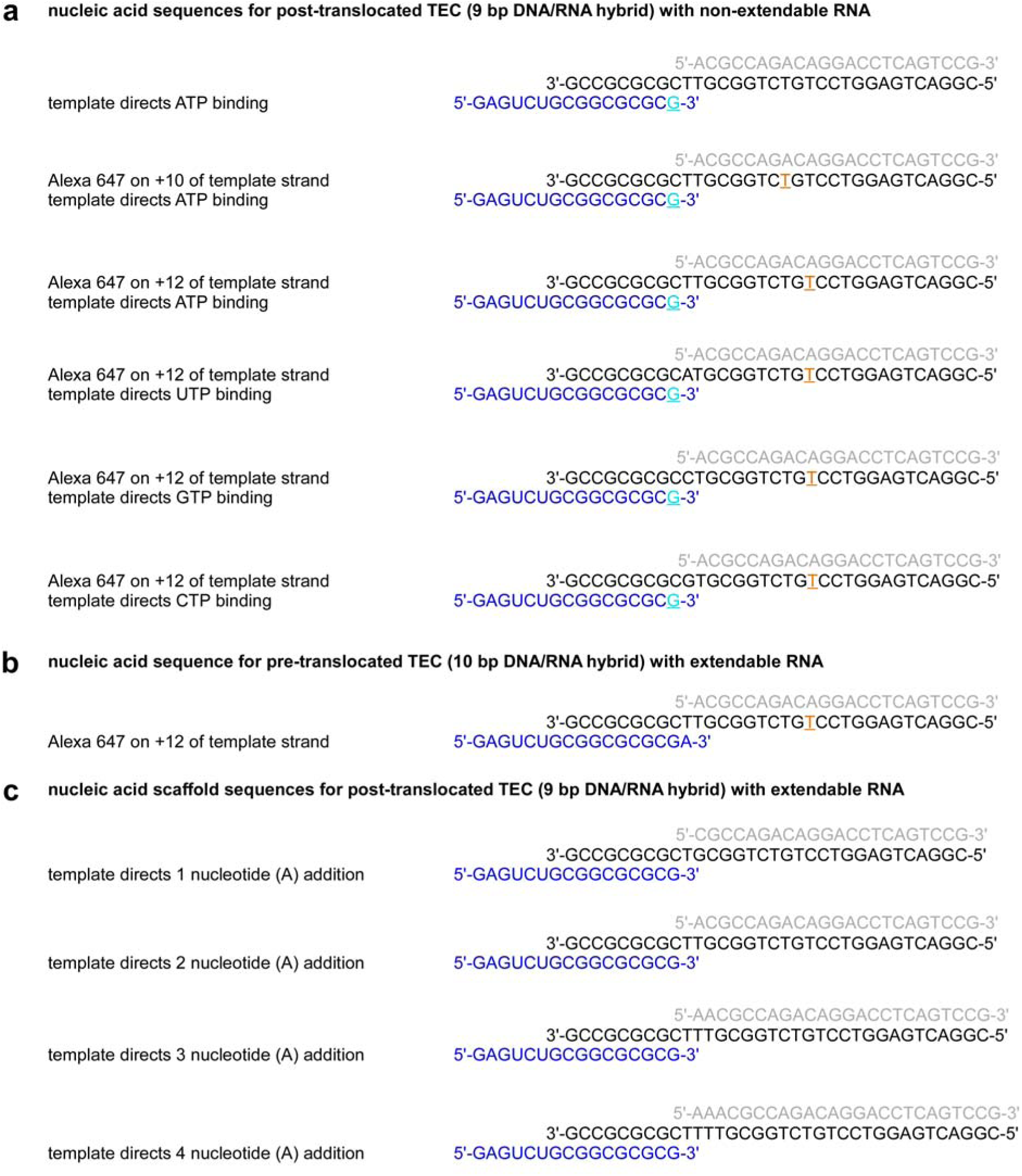
Nucleic-acid scaffolds. **a,** Nucleic-acid scaffolds for reconstitution of post-translocated TEC with 9 bp DNA-RNA hybrid with non-extendable RNA 3’ end (3’-deoxy). Black, DNA template strand; gray, DNA nontemplate strand; blue, RNA; cyan, non-extendable RNA 3’ end; orange, fluorescent-probe-labelled nucleotide. **b,** Nucleic-acid scaffold for reconstitution of pre-translocated TEC with 10 bp DNA-RNA hybrid. Black, DNA template strand; gray, DNA nontemplate strand; blue, RNA; orange, fluorescent-probe-labelled nucleotide. **c,** Nucleic-acid scaffolds for reconstitution of post-translocated TEC with 9 bp DNA-RNA hybrid and extendable RNA 3’ end (3’-hydroxy). Black, DNA template strand; gray, DNA nontemplate strand; blue, RNA.

**Fig. S2.**
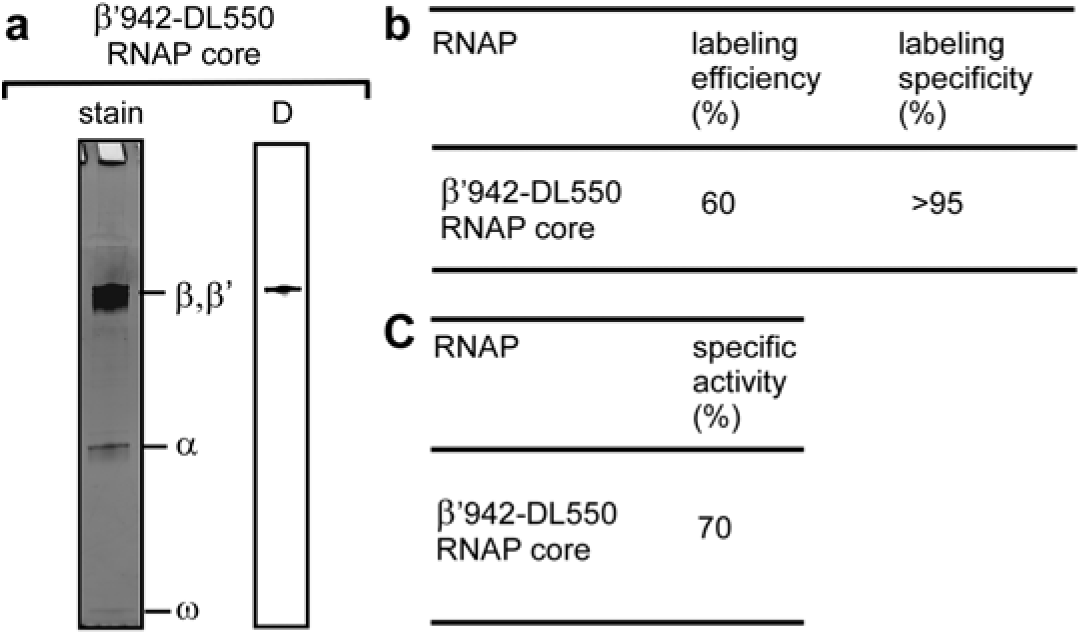
Incorporation of fluorescent probe into RNAP: singly-labelled RNAP derivative. **a,** Products of labelling reaction of RNAP derivative containing 4-azidophenylalanine (AzF) at position 942 of β’ subunit with Dylight 550 phosphine, as detected by Coomassie staining (left) and fluorescent scanning in donor-emission channel (532 nm excitation and 580 nm emission bandpass filters; right). Fluorescent labelling is observed only for the β’ subunit. **b,** Labelling efficiencies and labelling specificities (see Materials and Methods). **c,** Transcriptional activities of fluorescent-probe-labelled RNAP.

**Fig. S3.**
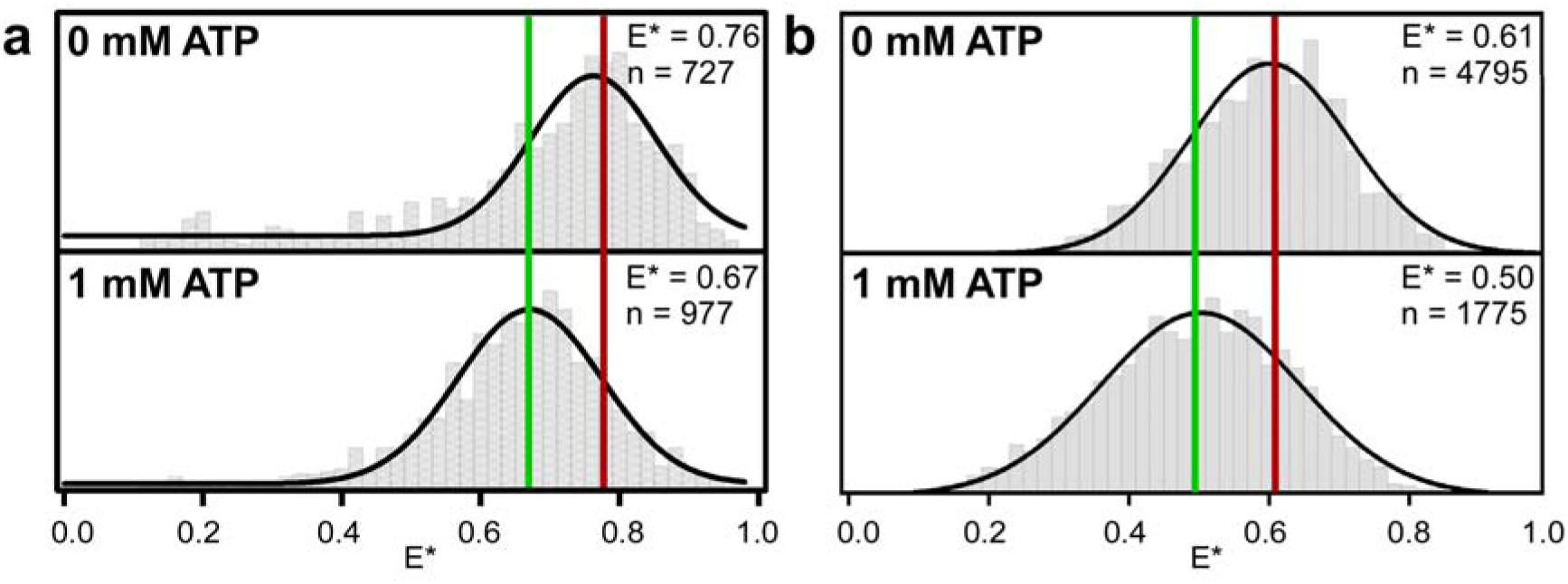
Use of smFRET to detect and characterize TL closing and opening in solution: confocal smFRET data. **a,** smFRET data for TEC in post-translocated state with a donor fluorescent probe in RNAP TL and acceptor fluorescent probe at position +10 of DNA template strand DNA of nucleic-acid scaffold. Data are shown for absence of NTP (top) and presence of saturating concentration of complementary NTP (1 mM ATP; bottom). Histogram and Gaussian fit of E* provide mean E* values for open-TL (red line) and closed-TL (green line) states. **b,** As a, but for TEC in post-translocated state with donor fluorescent probe in RNAP TL and acceptor fluorescent probe at position +12 of DNA template strand of nucleic-acid scaffold.

**Fig. S4.**
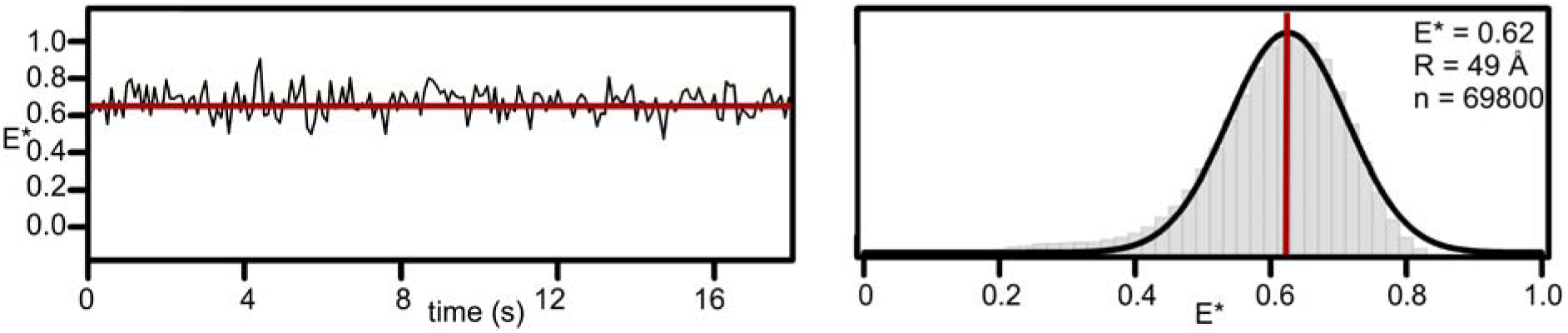
smFRET data for TEC in pre-translocated state in absence of NTP. Left subpanel, representative time traces of donor-acceptor FRET efficiency, E*, showing open-TL state (red). Right subpanel, histogram and Gaussian fit of E*, showing mean E* value of open-TL state (red line). R, mean donor-acceptor distance.

**Fig. S5.**
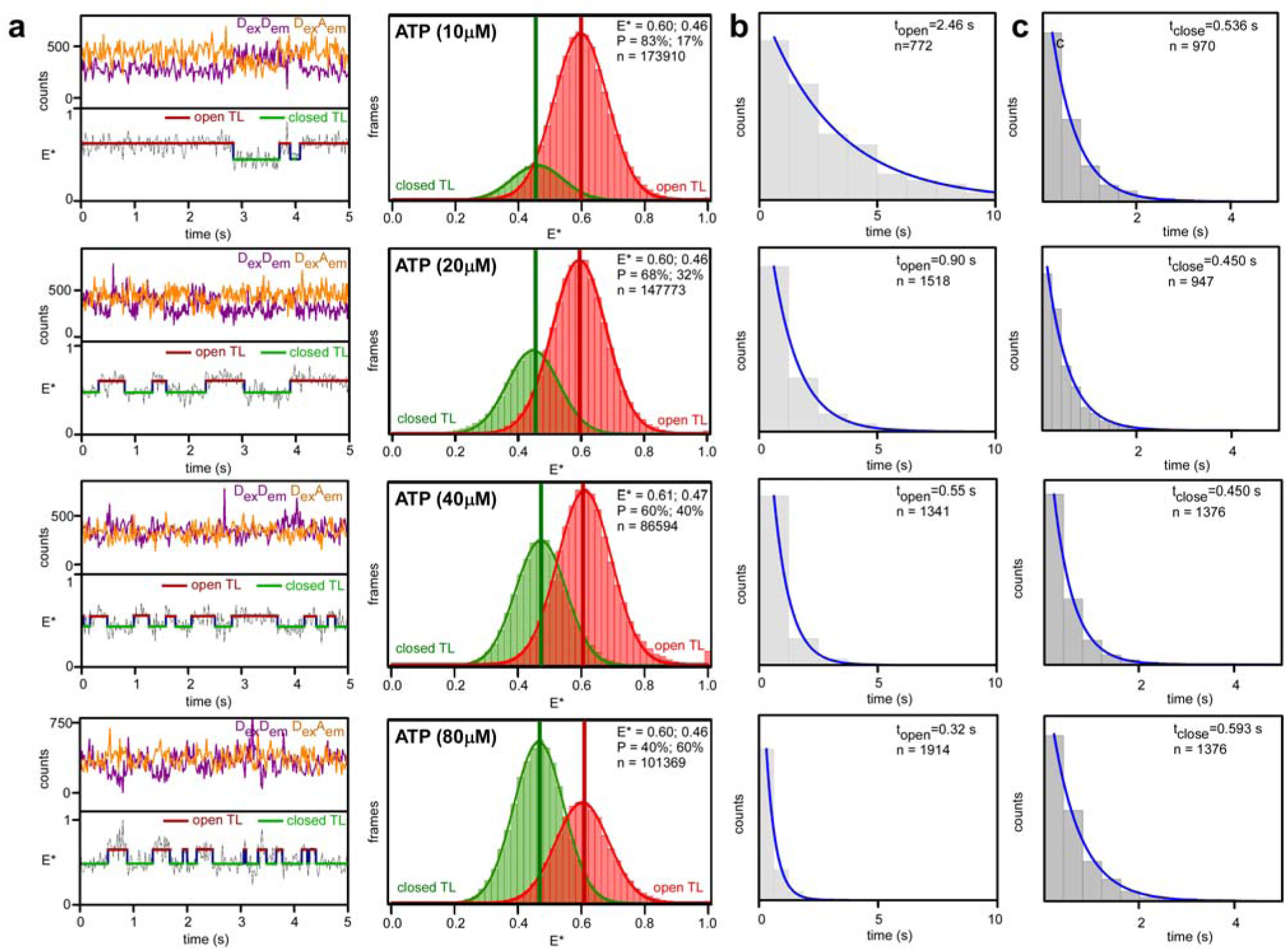
TL closing and opening occur on millisecond timescales: additional smFRET data in presence of sub-saturating concentrations of complementary NTP. **a,** smFRET data for TEC in post-translocated state in presence of each of four sub-saturating concentrations of complementary NTP (10, 20, 40, and 80 μM ATP). Left top, representative time trace of donor emission (purple) and acceptor emission (orange). Left, bottom, representative time trace of donor-acceptor FRET efficiency, E*, showing hidden-Markov-model (HMM)-assigned open-TL states (red), closed-TL states (green), and interstate transitions (blue). Right, histograms and Gaussian fits of E* show mean E* values of open-TL (red lines) and closed-TL (green lines) states. P, subpopulation percentage; R, mean donor-acceptor distance. **b,** Dwell-time distributions of open-TL states. **c,** Dwell-time distribution of closed-TL states.

**Fig. S6.**
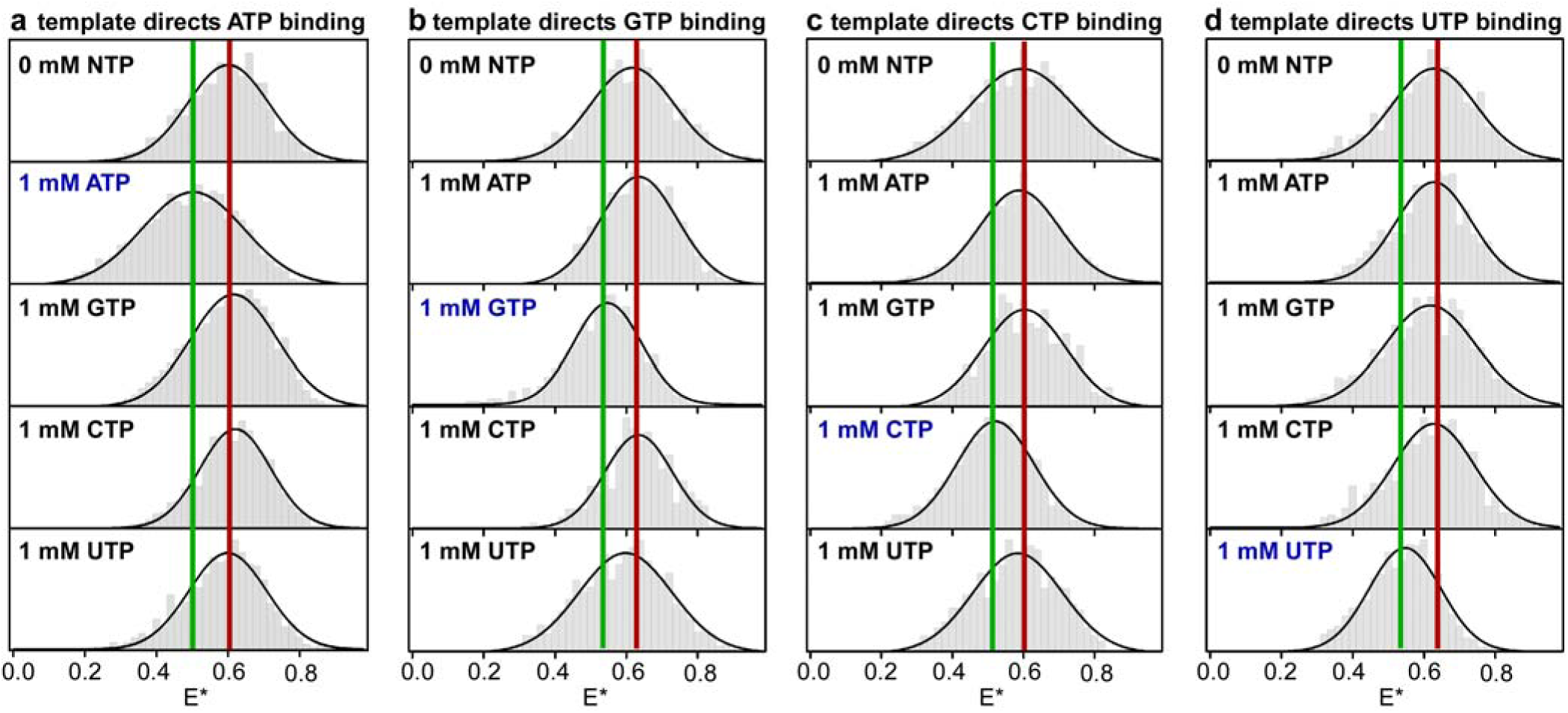
TL closing and opening can provide a checkpoint for NTP complementarity: additional smFRET data on effects of NTP complementarity on TL conformation. **a,** Effects of complementary and non-complementary NTPs on TL conformation with template directing binding of ATP. Histograms and Gaussian fits of E* show mean E* values of open-TL (red line) and closed-TL (green line) states. **b,** As a, but with template directing binding of GTP. **c,** As a, but with template directing binding of CTP. **d,** As a, but with template directing binding of UTP.

**Fig. S7.**
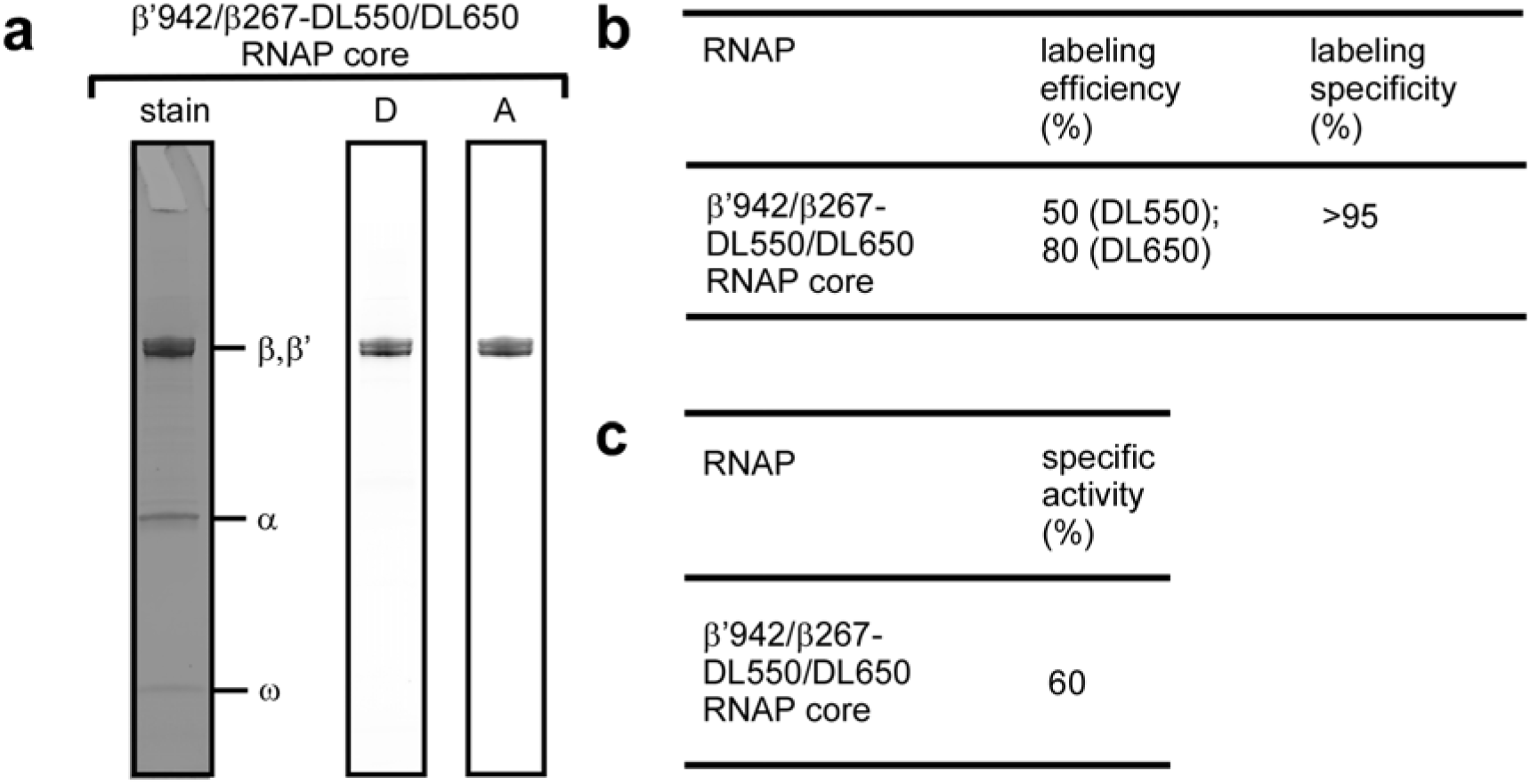
Incorporation of fluorescent probes into RNAP: doubly-labelled RNAP derivative. **a,** Products of stochastic labelling reaction of RNAP derivative containing 4-azidophenylalanine (AzF) at position 942 of β’ subunit and position 267 of β subunit with Dylight 550 phosphine and Dylight 650 phosphine, as detected by Coomassie staining (left), fluorescent scanning in donor-emission channel (D; 532 nm excitation and 580 nm emission bandpass filters; center) and fluorescent scanning in acceptor-emission channel (A; 633 nm excitation and 670 nm emission bandpass filters; right). Fluorescent labelling is observed only for β’ and β subunits. **b,** Labelling efficiencies and labelling specificities (see Materials and Methods). **c,** Transcriptional activities of fluorescent-probe-labelled RNAP.

**Fig. S8.**
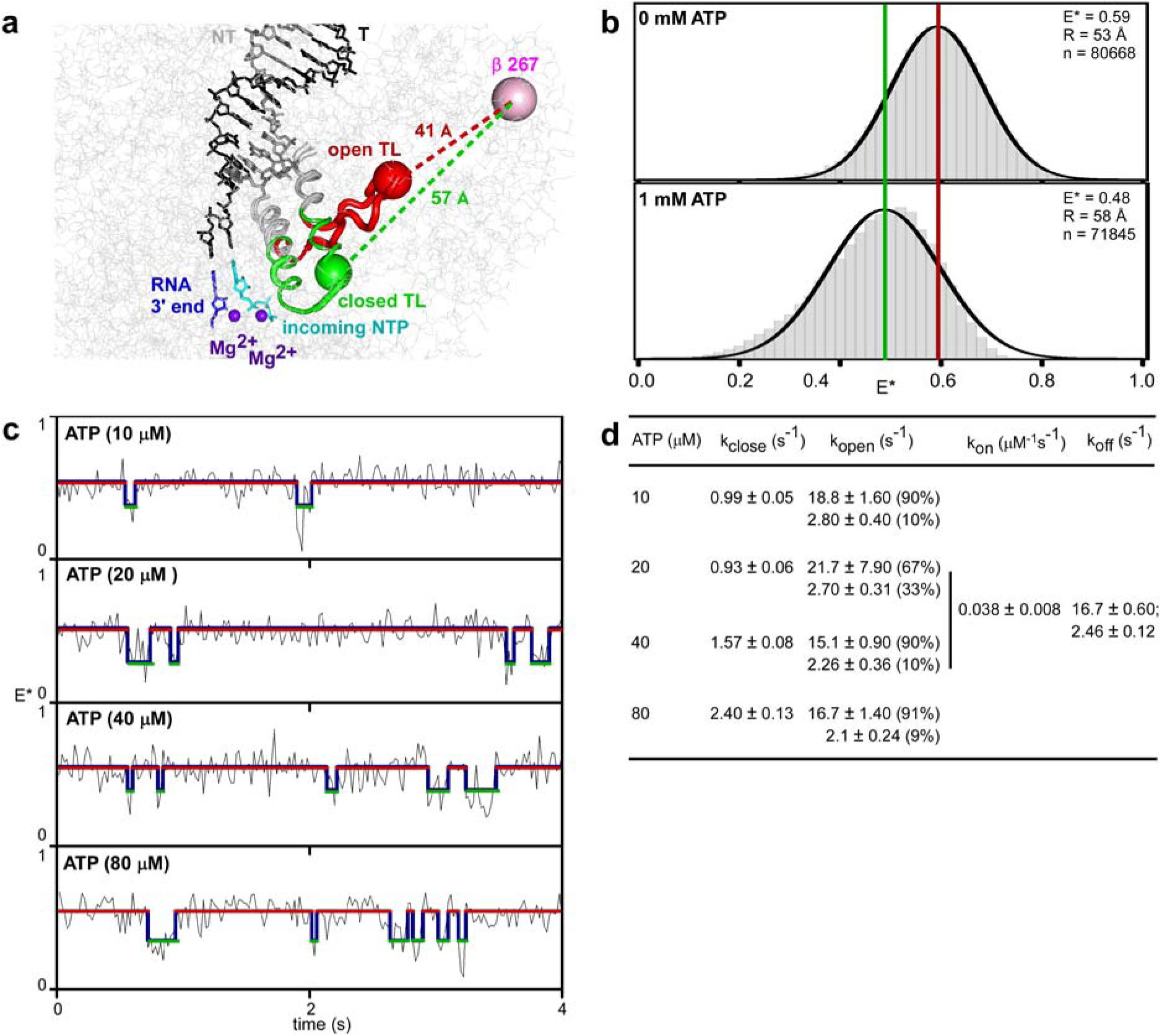
Use of smFRET to detect and characterize TL closing and opening in solution: doubly-labelled RNAP derivative. **a,** Measurement of smFRET between first fluorescent probe incorporated at tip of RNAP TL (red sphere for open-TL state; green sphere for closed-TL) and second fluorescent probe incorporated into position in RNAP β subunit (pink sphere). Predicted probe-probe distances are ~41 Å for open-TL state and ~57 Å for closed-TL state. Open-TL (red) and closed-TL (green) conformational states are as observed in crystal structures (*68, 69;* PDB 1ZYR and PDB 2O5J). RNAP core structure (gray lines) is as observed in (*70*; PDB 4YLN). Gray and red ribbon, RNAP trigger helices and TL in open-TL state; gray and green ribbon, RNAP trigger helices and TL in closed-TL state; gray and black sticks, DNA non-template and template strands; blue sticks, RNA 3’ nucleotide; cyan sticks, incoming NTP; purple spheres, catalytic Mg^2+^ ions Mg^2+^(I) and Mg^2+^(II). **b,** smFRET data for TEC in post-translocated state in absence of NTP (top) and in presence of saturating concentration of complementary NTP (1 mM ATP; bottom). Histograms and Gaussian fits of E* show mean E* values for open-TL (red line) and closed-TL (green line) states. R, mean donor-acceptor distance. **c,** smFRET data for TEC in post-translocated state in presence of each of four sub-saturating concentrations of complementary NTP (10, 20, 40, and 80 μM ATP). Representative time traces of donor-acceptor FRET efficiency, E*, showing hidden-Markov-model (HMM)-assigned open-TL states (red), closed-TL states (green), and interstate transitions (blue). **d,** TL-closing rate (k_close_), TL-opening rate (k_open_), ATP on-rate (k_on_), and ATP off-rate (k_off_) from experiments of a-c.

**Fig. S9.**
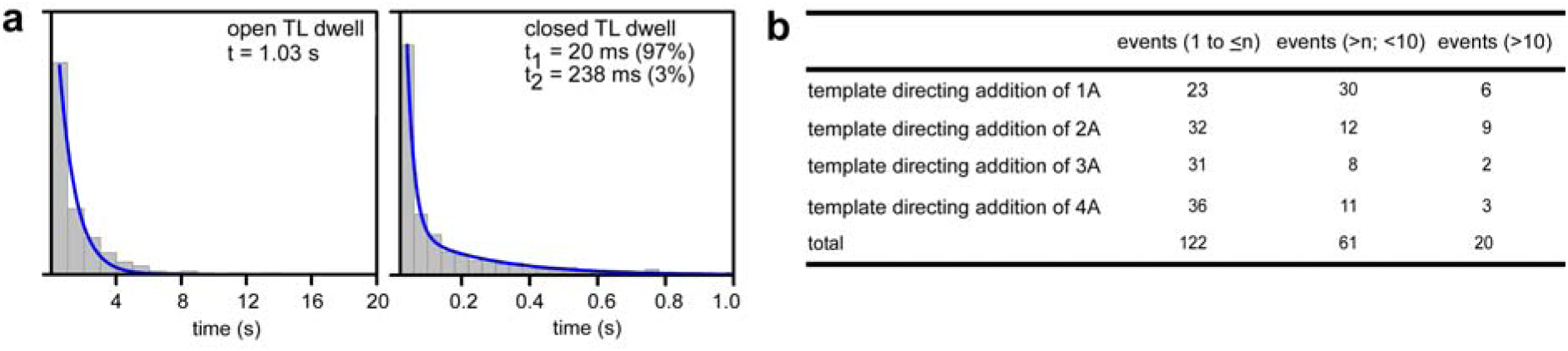
One TL closing-opening cycle typically occurs in each nucleotide addition in transcription elongation: TL-closing and TL-opening dynamics during nucleotide addition. **a,** Dwell-time distributions for open-TL (left) and closed-TL (right) states during nucleotide addition. **b,** Number of observed TL closing-opening events for templates directing 1, 2, 3, or 4 additions of A.

**Table S1:**
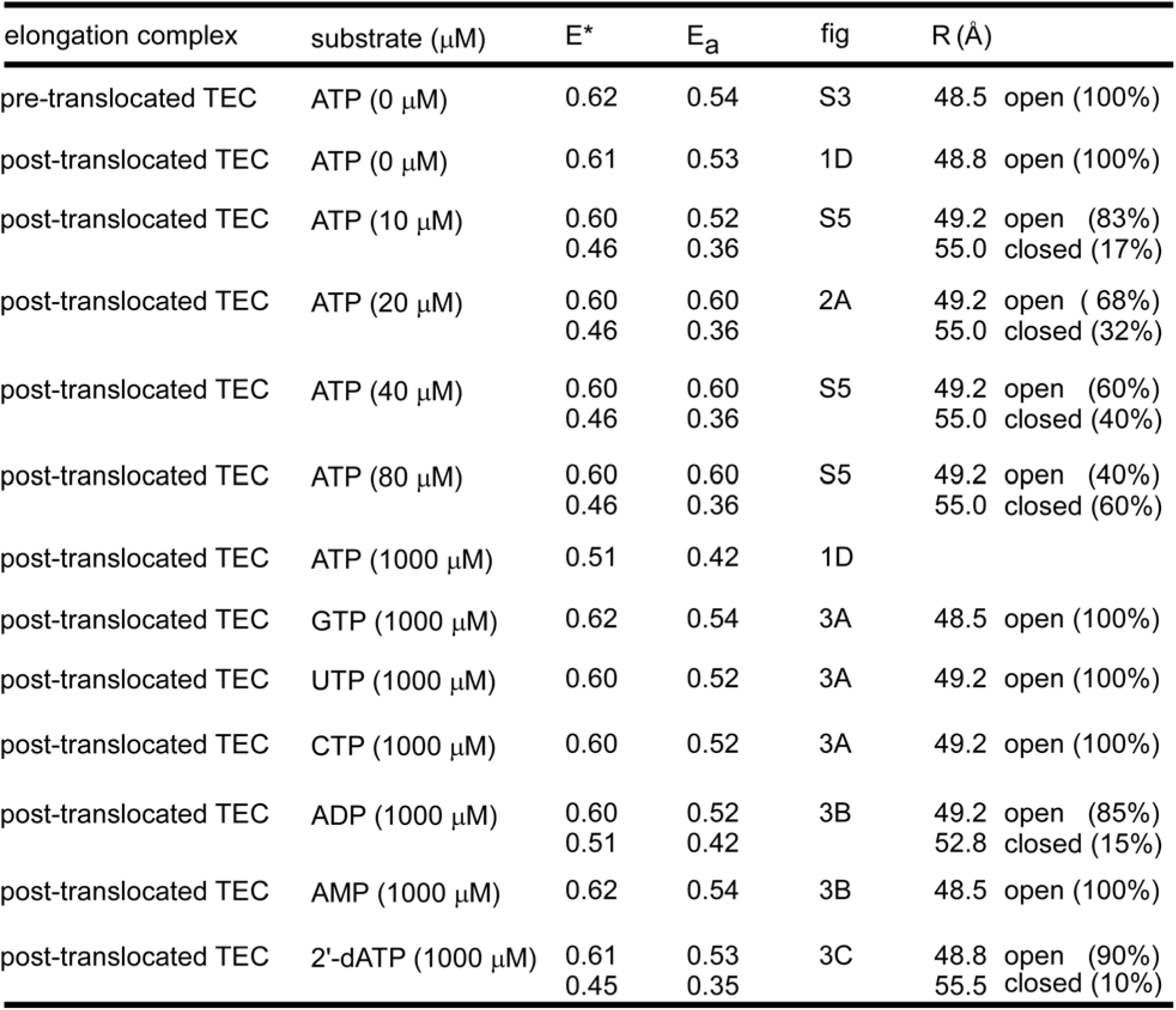
Apparent (E*) and accurate (E_a_) FRET efficiencies and distances (R): DL550 on tip of trigger loop (β’ residue 942) and Alexa 647 on +12 of the template strand

| elongation complex | substrate ($\mu\text{M}$) | $E^*$ | $E_a$ | fig | $R$ ($\text{\AA}$) |
| --- | --- | --- | --- | --- | --- |
| pre-translocated TEC | ATP (0 $\mu\text{M}$) | 0.62 | 0.54 | S3 | 48.5 open (100%) |
| post-translocated TEC | ATP (0 $\mu\text{M}$) | 0.61 | 0.53 | 1D | 48.8 open (100%) |
| post-translocated TEC | ATP (10 $\mu\text{M}$) | 0.60<br>0.46 | 0.52<br>0.36 | S5 | 49.2 open (83%)<br>55.0 closed (17%) |
| post-translocated TEC | ATP (20 $\mu\text{M}$) | 0.60<br>0.46 | 0.60<br>0.36 | 2A | 49.2 open (68%)<br>55.0 closed (32%) |
| post-translocated TEC | ATP (40 $\mu\text{M}$) | 0.60<br>0.46 | 0.60<br>0.36 | S5 | 49.2 open (60%)<br>55.0 closed (40%) |
| post-translocated TEC | ATP (80 $\mu\text{M}$) | 0.60<br>0.46 | 0.60<br>0.36 | S5 | 49.2 open (40%)<br>55.0 closed (60%) |
| post-translocated TEC | ATP (1000 $\mu\text{M}$) | 0.51 | 0.42 | 1D | |
| post-translocated TEC | GTP (1000 $\mu\text{M}$) | 0.62 | 0.54 | 3A | 48.5 open (100%) |
| post-translocated TEC | UTP (1000 $\mu\text{M}$) | 0.60 | 0.52 | 3A | 49.2 open (100%) |
| post-translocated TEC | CTP (1000 $\mu\text{M}$) | 0.60 | 0.52 | 3A | 49.2 open (100%) |
| post-translocated TEC | ADP (1000 $\mu\text{M}$) | 0.60<br>0.51 | 0.52<br>0.42 | 3B | 49.2 open (85%)<br>52.8 closed (15%) |
| post-translocated TEC | AMP (1000 $\mu\text{M}$) | 0.62 | 0.54 | 3B | 48.5 open (100%) |
| post-translocated TEC | 2'-dATP (1000 $\mu\text{M}$) | 0.61<br>0.45 | 0.53<br>0.35 | 3C | 48.8 open (90%)<br>55.5 closed (10%) |

**Table S2:** Apparent (E*) and accurate (E_a_) FRET efficiencies and distances (R): DL550/DL650 on tip of trigger loop (β’ residue 942) and β subunit residue 267

| elongation complex | substrate ($\mu\text{M}$) | $E^*$ | $E_a$ | fig | R ($\text{\AA}$) | |
| --- | --- | --- | --- | --- | --- | --- |
| post-translocated TEC | ATP (0 $\mu\text{M}$) | 0.59 | 0.46 | S8 | 53 | open (100%) |
| post-translocated TEC | ATP (1000 $\mu\text{M}$) | 0.48 | 0.34 | S8 | 58 | closed (100%) |

## References

1. A. Mazumder, A.N. Kapanidis, Recent advances in understanding σ^70^-dependent transcription initiation mechanisms. J. Mol. Biol., in press (2019).

2. G.A. Belogurov, I. Artsimovitch, The mechanisms of substrate selection, catalysis, and translocation by the elongating RNA polymerase. J. Mol. Biol., in press (2019).

3. J.W. Roberts, Mechanisms of bacterial transcription termination. J. Mol. Biol., in press (2019).

4. E.A. Abbondanzieri, W.J. Greenleaf, J.W. Shaevitz, R. Landick, S.M. Block, Direct observation of base-pair stepping by RNA polymerase. Nature 438, 460–465 (2005).

5. L.R. Zhang and R. Landick, Substrate loading, nucleotide addition, and translocation by RNA polymerase, In RNA Polymerases as Molecular Motors, Eds. Buc, H., Strick, S., Royal Society of Chemistry, London (2009).

6. M.H. Larson, R. Landick, S.M. Block, Single-molecule studies of RNA polymerase: one singular sensation, every little step it takes. Mol. Cell 41, 249–262 (2011).

7. D.A. Erie, T.D. Yager, P.H. von Hippel, The single-nucleotide addition cycle in transcription: a biophysical and biochemical perspective. Annu. Rev. Biophys. Biomol. Struct. 21, 379–415 (1992).

8. D. Wang, D.A. Bushnell, K.D. Westover, C.D. Kaplan, R.D. Kornberg, Structural basis of transcription: role of the trigger loop in substrate specificity and catalysis. Cell 127, 941–954 (2006).

9. D.G. Vassylyev, M.N. Vassylyeva, J. Zhang, M. Palangat, I. Artsimovitch, R. Landick, Structural basis for substrate loading in bacterial RNA polymerase. Nature 448, 163–168 (2007).

10. F. Brueckner, J. Ortiz, P. Cramer, A movie of the RNA polymerase nucleotide addition cycle. Curr. Opin. Struct. Biol. 19, 294–299 (2009).

11. F.W. Martinez-Rucobo, P. Cramer, Structural basis of transcription elongation. Biochim. Biophys. Acta. 1829, 9–19 (2013).

12. I. Toulokhonov, J. Zhang, M. Palangat, R. Landick, A central role of the RNA polymerase trigger loop in active-site rearrangement during transcriptional pausing. Mol. Cell 27, 406–419 (2007).

13. F. Brueckner, P. Cramer, Structural basis of transcription inhibition by alpha-amanitin and implications for RNA polymerase II translocation. Nat. Struct. Mol. Biol. 15, 811–818 (2008).

14. C.D. Kaplan, K.M. Larsson, R.D. Kornberg, The RNA polymerase II trigger loop functions in substrate selection and is directly targeted by alpha-amanitin. Mol. Cell 30, 547–556 (2008).

15. L. Tan, S. Wiesler, D. Trzaska, H.C. Carney, R.O. Weinzierl, Bridge helix and trigger loop perturbations generate superactive RNA polymerases. J. Biol. 7, 40 (2008).

16. J. Zhang, M. Palangat, R. Landick, Role of the RNA polymerase trigger loop in catalysis and pausing. Nat. Struct. Mol. Biol. 17, 99–104 (2010).

17. X. Huang, D. Wang, D.R. Weiss, D.A. Bushnell, R.D. Kornberg, M. Levitt, RNA polymerase II trigger loop residues stabilize and position the incoming nucleotide triphosphate in transcription. Proc. Natl. Acad. Sci. U S A 107, 15745–15750 (2010).

18. M.H. Larson, J. Zhou, C.D. Kaplan, M. Palangat, R.D. Kornberg, R. Landick, S.M. Block, Trigger loop dynamics mediate the balance between the transcriptional fidelity and speed of RNA polymerase II. Proc. Natl. Acad. Sci. U S A 109, 6555–6560 (2012).

19. C.D. Kaplan, H. Jin, I.L. Zhang, A. Belyanin, Dissection of Pol II trigger loop function and Pol II activity-dependent control of start site selection *in vivo*. PLoS. Genet. 8, e1002627 (2012).

20. T. Fouqueau, M.E. Zeller, A.C. Cheung, P. Cramer, M. Thomm, The RNA polymerase trigger loop functions in all three phases of the transcription cycle. Nucleic Acids Res. 41, 7048–7059 (2013).

21. B. Wang, A.V. Predeus, Z.F. Burton, M. Feig, Energetic and structural details of the trigger-loop closing transition in RNA polymerase II. Biophys. J. 105, 767–775 (2013).

22. D. Nayak, M. Voss, T. Windgassen, R.A. Mooney, R. Landick, Cys-pair reporters detect a constrained trigger loop in a paused RNA polymerase. Mol. Cell 50, 882–893 (2013).

23. N. Miropolskaya, D. Esyunina, S. Klimasauskas, V. Nikiforov, I. Artsimovitch, A. Kulbachinskiy, Interplay between the trigger loop and the F loop during RNA polymerase catalysis. Nucleic Acids Res. 42, 544–552 (2014).

24. L. Xu, K.V. Butler, J. Chong, J. Wengel, E.T. Kool, D. Wang, Dissecting the chemical interactions and substrate structural signatures governing RNA polymerase II trigger loop closure by synthetic nucleic acid analogues. Nucleic Acids Res. 42, 5863–5870 (2014).

25. T.A. Windgassen, R.A. Mooney, D. Nayak, M. Palangat, J. Zhang, R. Landick, Trigger-helix folding pathway and SI3 mediate catalysis and hairpin-stabilized pausing by *Escherichia coli* RNA polymerase. Nucleic Acids Res. 42, 12707–12721 (2014).

26. Y.X. Mejia, E. Nudler, C. Bustamante, Trigger loop folding determines transcription rate of Escherichia coli’s RNA polymerase. Proc. Natl. Acad. Sci. U S A 112, 743–748 (2015).

27. C. Qiu, O.C. Erinne, J.M. Dave, P. Cui, H. Jin, N. Muthukrishnan, L.K. Tang, S.G. Babu, K.C. Lam, P. J. Vandeventer, R. Strohner, J. Van den Brulle, S.H. Sze, C.D. Kaplan, High-resolution phenotypic landscape of the RNA polymerase II trigger loop. PLoS. Genet. 12, e1006321 (2016).

28. T.V. Mishanina, M.Z. Palo, D. Nayak, R.A. Mooney, R. Landick, Trigger loop of RNA polymerase is a positional, not acid-base, catalyst for both transcription and proofreading. Proc. Natl. Acad. Sci. U S A 114, E5103–E5112 (2017).

29. S. Tuske, S.G. Sarafianos, X. Wang, B. Hudson, E. Sineva, J. Mukhopadhyay, J.J. Birktoft, O. Leroy, S. Ismail, A.D. Clark, Jr., C. Dharia, A. Napoli, O. Laptenko, J. Lee, S. Borukhov, R.H. Ebright, E. Arnold, Inhibition of bacterial RNA polymerase by streptolydigin: stabilization of a straight-bridge-helix active-center conformation. Cell 122, 541–552 (2005).

30. D. Temiakov, N. Zenkin, M.N. Vassylyeva, A. Perederina, T.H. Tahirov, E. Kashkina, M. Savkina, S. Zorov, V. Nikiforov, N. Igarashi, N. Matsugaki, S. Wakatsuki, K. Severinov, D.G. Vassylyev, Structural basis of transcription inhibition by antibiotic streptolydigin. Mol. Cell 19, 655–666 (2005).

31. D. Degen, Y. Feng, Y. Zhang, K.Y. Ebright, Y.W. Ebright, M. Gigliotti, H. Vahedian-Movahed, S. Mandal, M. Talaue, N. Connell, E. Arnold, W. Fenical, R.H. Ebright, Transcription inhibition by the depsipeptide antibiotic salinamide A. Elife 3, e02451 (2014).

32. Y. Feng, D. Degen, X. Wang, M. Gigliotti, S. Liu, Y. Zhang, D. Das, T. Michalchuk, Y.W. Ebright, M. Talaue, N. Connell, R.H. Ebright, Structural basis of transcription inhibition by CBR hydroxamidines and CBR pyrazoles. Structure 23, 1470–1481 (2015).

33. B. Bae, D. Nayak, A. Ray, A. Mustaev, R. Landick, S. A. Darst, CBR antimicrobials inhibit RNA polymerase via at least two bridge-helix cap-mediated effects on nucleotide addition. Proc. Natl. Acad. Sci. U S A 112, E4178–4187 (2015).

34. W. Lin, S. Mandal, D. Degen, Y. Liu, Y.W. Ebright, S. Li, Y. Feng, Y. Zhang, S. Mandal, Y. Jiang, S. Liu, M. Gigliotti, M. Talaue, N. Connell, K. Das, E. Arnold, R.H. Ebright, Structural basis of *Mycobacterium tuberculosis* transcription and transcription inhibition. Mol. Cell 66, 169–179 e168 (2017).

35. R.H. Ebright, Y.W. Ebright, S. Mandal, R. Wilde, and S. Li, Antibacterial agents: N(alpha)-aroyl-N-aryl-phenylalaninamides. US Patent US9919998.36 (2018).

36. N.R. Braffman, F.J. Piscotta, J. Hauver, E.A. Campbell, A.J. Link, S.A. Darst, Structural mechanism of transcription inhibition by lasso peptides microcin J25 and capistruin. Proc. Natl. Acad. Sci. U S A 116, 1273–1278 (2019).

37. A. Chakraborty, D. Wang, Y.W. Ebright, Y. Korlann, E. Kortkhonjia, T. Kim, S. Chowdhury, S. Wigneshweraraj, H. Irschik, R. Jansen, B.T. Nixon, J. Knight, S. Weiss, R.H. Ebright, Opening and closing of the bacterial RNA polymerase clamp. Science 337, 591–595 (2012).

38. A. Chakraborty, A. Mazumder, M. Lin, A. Hasemeyer, Q. Xu, D. Wang, Y.W. Ebright, R.H. Ebright, Site-specific incorporation of probes into RNA polymerase by unnatural-amino-acid mutagenesis and Staudinger-Bertozzi ligation. Methods Mol. Biol. 1276, 101–131 (2015).

39. D. Duchi, A. Mazumder, A.M. Malinen, R.H. Ebright, A.N. Kapanidis, The RNA polymerase clamp interconverts dynamically among three states and is stabilized in a partly closed state by ppGpp. Nucleic Acids Res. 46, 7284–7295 (2018).

40. W. Lin, K. Das, D. Degen, A. Mazumder, D. Duchi, D. Wang, Y.W. Ebright, R.Y. Ebright, E. Sineva, M. Gigliotti, A. Srivastava, S. Mandal, Y. Jiang, Y. Liu, R. Yin, Z. Zhang, E.T. Eng, D. Thomas, S. Donadio, H. Zhang, C. Zhang, A.N. Kapanidis, R.H. Ebright, Structural basis of transcription inhibition by fidaxomicin (lipiarmycin A3). Mol. Cell 70, 60–71 e15 (2018).

41. M. Chlenov, S. Masuda, K. S. Murakami, V. Nikiforov, S. A. Darst, A. Mustaev, Structure and function of lineage-specific sequence insertions in the bacterial RNA polymerase beta’ subunit. J Mol Biol 353, 138–154 (2005).

42. A.N. Kapanidis, N.K. Lee, T.A. Laurence, S. Doose, E. Margeat, S. Weiss, Fluorescence-aided molecule sorting: analysis of structure and interactions by alternating-laser excitation of single molecules. Proc. Natl. Acad. Sci. U S A 101, 8936–8941 (2004).

43. S.J. Holden, S. Uphoff, J. Hohlbein, D. Yadin, L. Le Reste, O.J. Britton, A.N. Kapanidis, Defining the limits of single-molecule FRET resolution in TIRF microscopy. Biophys. J. 99, 3102–3111 (2010).

44. E. Margeat, A.N. Kapanidis, P. Tinnefeld, Y. Wang, J. Mukhopadhyay, R.H. Ebright, S. Weiss, Direct observation of abortive initiation and promoter escape within single immobilized transcription complexes. Biophys J 90, 1419–1431 (2006).

45. D. Duchi, D.L. Bauer, L. Fernandez, G. Evans, N. Robb, L.C. Hwang, K. Gryte, A. Tomescu, P. Zawadzki, Z. Morichaud, K. Brodolin, A.N. Kapanidis, RNA polymerase pausing during initial transcription. Mol Cell 63, 939–950 (2016).

46. R.S. Johnson, M. Strausbauch, J.K. Carraway, Rapid pyrophosphate release from transcriptional elongation complexes appears to be coupled to a nucleotide-induced conformational change in *E. coli* core polymerase. J. Mol. Biol. 412, 849–861 (2011).

47. E.A. Campbell, N. Korzheva, A. Mustaev, K. Murakami, S. Nair, A. Goldfarb, S.A. Darst, Structural mechanism for rifampicin inhibition of bacterial RNA polymerase. Cell 104, 901–912 (2001).

48. I.M. Derrington, J.M. Craig, E. Stava, A.H. Laszlo, B.C. Ross, H. Brinkerhoff, I.C. Nova, K. Doering, B.I. Tickman, M. Ronaghi, J.G. Mandell, K.L. Gunderson, J.H. Gundlach, Subangstrom single-molecule measurements of motor proteins using a nanopore. Nat. Biotechnol. 33, 1073–1075 (2015).

## Supplementary References

49. J.W. Chin, S.W. Santoro, A.B. Martin, D.S. King, L. Wang, P.G. Schultz, Addition of p-azido-L-phenylalanine to the genetic code of Escherichia coli. J Am Chem Soc 124, 9026–9027 (2002).

50. A. Chakraborty, D. Wang, Y.W. Ebright, Y. Korlann, E. Kortkhonjia, T. Kim, S. Chowdhury, S. Wigneshweraraj, H. Irschik, R. Jansen, B.T. Nixon, J. Knight, S. Weiss, R.H. Ebright, Opening and closing of the bacterial RNA polymerase clamp. Science 337, 591–595 (2012).

51. D. Duchi, A. Mazumder, A.M. Malinen, R.H. Ebright, A.N. Kapanidis, The RNA polymerase clamp interconverts dynamically among three states and is stabilized in a partly closed state by ppGpp. Nucleic Acids Res 46, 7284–7295 (2018).

52. W. Lin, K. Das, D. Degen, A. Mazumder, D. Duchi, D. Wang, Y.W. Ebright, R.Y. Ebright, E. Sineva, M. Gigliotti, A. Srivastava, S. Mandal, Y. Jiang, Y. Liu, R. Yin, Z. Zhang, E.T. Eng, D. Thomas, S. Donadio, H. Zhang, C. Zhang, A.N. Kapanidis, R.H. Ebright, Structural basis of transcription inhibition by Fidaxomicin (Lipiarmycin A3). Mol Cell 70, 60–71 e15 (2018).

53. B.P. Hudson, J. Quispe, S. Lara-Gonzalez, Y. Kim, H.M. Berman, E. Arnold, R.H. Ebright, C.L. Lawson, Three-dimensional EM structure of an intact activator-dependent transcription initiation complex. Proc Natl Acad Sci U S A 106, 19830–19835 (2009).

54. C.E. Vrentas, T. Gaal, W. Ross, R.H. Ebright, R.L. Gourse, Response of RNA polymerase to ppGpp: requirement for the omega subunit and relief of this requirement by DksA. Genes Dev 19, 2378–2387 (2005).

55. J. Sambrook, D. Russell, Molecular cloning: A laboratory manual (Cold Spring harbour, NY, Cold Spring Haror Laboratory) (2001).

56. W. Niu, Y. Kim, G. Tau, T. Heyduk, R.H. Ebright, Transcription activation at class II CAP-dependent promoters: two interactions between CAP and RNA polymerase. Cell 87, 1123–1134 (1996).

57. D. Degen, Y. Feng, Y. Zhang, K.Y. Ebright, Y.W. Ebright, M. Gigliotti, H. Vahedian-Movahed, S. Mandal, M. Talaue, N. Connell, E. Arnold, W. Fenical, R.H. Ebright, Transcription inhibition by the depsipeptide antibiotic salinamide A. eLife 3, e02451 (2014).

58. Y. Zhang, D. Degen, M.X. Ho, E. Sineva, K.Y. Ebright, Y.W. Ebright, V. Mekler, H. Vahedian-Movahed, Y. Feng, R. Yin, S. Tuske, H. Irschik, R. Jansen, S. Maffioli, S. Donadio, E. Arnold, R.H. Ebright, GE23077 binds to the RNA polymerase ‘i’ and ‘i+1’ sites and prevents the binding of initiating nucleotides. eLife 3, e02450 (2014).

59. A.N. Kapanidis, N.K. Lee, T.A. Laurence, S. Doose, E. Margeat, S. Weiss, Fluorescence-aided molecule sorting: analysis of structure and interactions by alternating-laser excitation of single molecules. Proc Natl Acad Sci U S A 101, 8936–8941 (2004).

60. J. Mukhopadhyay, E. Sineva, J. Knight, R.M. Levy, R.H. Ebright, Antibacterial peptide microcin J25 inhibits transcription by binding within and obstructing the RNA polymerase secondary channel. Mol Cell 14, 739–751 (2004).

61. Ebright, R., Ebright, Y., Mandal, S., Wilde, R., and Li, Shengjian. (2018) Antibacterial agents: N(alpha)-aroyl-N-aryl-phenylalaninamides. US Patent US9919998.36.

62. N.K. Lee, A.N. Kapanidis, Y. Wang, X. Michalet, J. Mukhopadhyay, R.H. Ebright, S. Weiss, Accurate FRET measurements within single diffusing biomolecules using alternating-laser excitation. Biophys J 88, 2939–2953 (2005).

63. S.J. Holden, S. Uphoff, J. Hohlbein, D. Yadin, L. Le Reste, O.J. Britton, A.N. Kapanidis, Defining the limits of single-molecule FRET resolution in TIRF microscopy. Biophys J 99, 3102–3111 (2010).

64. D. Axelrod, N.L. Thompson, T.P. Burghardt, Total internal inflection fluorescent microscopy. J Microsc 129, 19–28 (1983).

65. J.W. van de Meent, J.E. Bronson, C.H. Wiggins, R.L. Gonzalez, Jr., Empirical Bayes methods enable advanced population-level analyses of single-molecule FRET experiments. Biophys J 106, 1327–1337 (2014).

66. S. Schulz, A. Gietl, K. Smollett, P. Tinnefeld, F. Werner, D. Grohmann, TFE and Spt4/5 open and close the RNA polymerase clamp during the transcription cycle. Proc Natl Acad Sci U S A 113, E1816–1825 (2016).

67. P. Wu, L. Brand, Orientation factor in steady-state and time-resolved resonance energy transfer measurements. Biochemistry 31, 7939–7947 (1992).

68. S. Tuske, S. G. Sarafianos, X. Wang, B. Hudson, E. Sineva, J. Mukhopadhyay, J. J. Birktoft, O. Leroy, S. Ismail, A. D. Clark, Jr., C. Dharia, A. Napoli, O. Laptenko, J. Lee, S. Borukhov, R. H. Ebright, E. Arnold, Inhibition of bacterial RNA polymerase by streptolydigin: stabilization of a straight-bridge-helix active-center conformation. Cell 122, 541–552 (2005).

69. D. G. Vassylyev, M. N. Vassylyeva, J. Zhang, M. Palangat, I. Artsimovitch, R. Landick, Structural basis for substrate loading in bacterial RNA polymerase. Nature 448, 163–168 (2007).

70. Y. Zuo, T.A. Steitz, Crystal structures of the E. coli transcription initiation complexes with a complete bubble. Mol Cell 58, 534–540 (2015).

